# Implementation and calibration of spectroscopic single-molecule localization microscopy

**DOI:** 10.1101/2024.08.15.608177

**Authors:** Benjamin Brenner, Wei-Hong Yeo, Youngseop Lee, Junghun Kweon, Cheng Sun, Hao F. Zhang

## Abstract

Spectroscopic single-molecule localization microscopy (sSMLM) allows multi-color super-resolution images with high spectral sensitivity. In this protocol, we provide essential information for researchers to implement sSMLM in a laboratory setting. We describe how to assemble and align the illumination and detection paths of a 3D dual-wedge prism (DWP)-based sSMLM instrument. We provide detailed step-by-step instructions for performing spectral and axial calibration using fluorescent beads and a nanohole array, respectively. We also discuss using sSMLM to image fluorescently labeled cells and report a new MATLAB package, RainbowSTORM v2, to reconstruct super-resolution 3D images. Further, we present representative images as typical anticipated results for users to validate.

## Introduction

Single-molecule localization microscopy (SMLM) relies on the computational localization of sparse fluorescent emitters to achieve super-resolution imaging [1–4]. In each frame of an SMLM dataset, only a small, random subset of labeled proteins are detected, which is generally achieved via blinking fluorophores [5], though in point accumulation for imaging in nanoscale topography or PAINT, free-floating fluorophores stochastically bind and unbind to the targeted protein [6]. Each single-molecule fluorescence emission forms a diffraction-limited point spread function (PSF) on the array detector in the shape of an airy disk pattern, often approximated by 2D Gaussian function. The true location of the single-molecule emitter for each PSF can be estimated by fitting the 2D Gaussian function [7]. Hence, the final SMLM image consists of all the identified localizations.

Spectroscopic SMLM or sSMLM extends the capabilities of SMLM by simultaneously capturing the spectral signatures of single molecular emissions, and has been used for multi-color imaging [1, 8], polarity sensing [9, 10], and chemical characterization [11, 12]. It adds a dispersive element (such as a prism or a diffraction grating) into the microscope’s imaging path, splitting it into a spatial channel and a spectral channel. Several groups introduced different implementations of sSMLM [13], including single-objective prism-based sSMLM [9], dual-objective prism-based sSMLM [1], and grating-based sSMLM [2]. Our group recently introduced dual wedge prism (DWP)-based sSMLM [14, 15], in which the beam-splitting and dispersive elements are compacted into a single, monolithic assembly with only two air-glass interfaces to minimize photon loss. The DWP assembly can also fit into the microscope side port for easy integration.

This protocol paper focuses on instrumentation, calibration, and image processing of sSMLM to help researchers integrate sSMLM into their work and promote standardized operation. Due to the different investigative needs of the samples, we can not focus on sample preparation in this protocol. While we will describe our protocol for fixing and labeling cells, this is meant as an example, not a comprehensive guidance. The procedures described here are compatible with a wide range of SMLM tagging methods, including immunolabeling [16], tags [17, 18], PAINT [6, 19], tracking [20–22], autofluorescence [23], organelle-targeting fluorophores [24], intrinsic fluorophore expression [25], and others of which the authors may not be aware.

While most of the setup and calibration proce-dures discussed here are adaptable to any implementa-tion of sSMLM, by way of example, the procedures section will focus on DWP-based sSMLM, which we consider to be the simplest 3D imaging implementation to set up and calibrate. DWP-based sSMLM is also less lossy than all other implementations except 4Pi sSMLM, which is more complex and imposes limitations on the sample size [13]. We will discuss the procedures for alignment, calibrating axial localization and spectral classification, image acquisition, and image analysis. For image analysis, our lab previously introduced an ImageJ plug-in RainbowSTORM 1.0 [26]. Here, we provide an updated and editable MATLAB package RainbowSTORM v2, which integrates the latest spectral calibration and classification methods and enables 3D imaging via DWP sSMLM. Users who wish to tailor RainbowSTORM for specific applications can revise our Matlab codes. We will also, by way of example, discuss the culture and labeling of cells for 3D multi-color imaging. However, protocols will vary for different biological samples. We will begin with a general overview of each procedure, followed by a detailed, step-by-step guide. We list the catalog number and source company of all the mentioned commercial components, supplies, and equipment in the materials section. For more specific information concerning the design parameters for DWP, readers can refer to the recent work by Song et al. [14, 15]

### Conceptual overview of sSMLM and calibration procedures

Each frame of sSMLM raw data consists of sparse, spatial PSFs on one side, and the corresponding spectra convolved with each spatial PSF on the other. We refer to them as the spatial PSFs and spectral PSFs, respectively. To spectrally classify each fluorescent emission, we pair the spatial PSF to its corresponding spectral PSF and then identify the centroids of both PSFs using Gaussian fitting. We then calculate the distance between the pair of centroids, and map this value to the to the peak emission spectrum, as fluorophore spectra with longer wavelengths will be right-shifted relative to their spatial PSF position. Before sample imaging, users must perform spectral calibration to determine how the centroid pair distance is mapped onto the spectral image calibrated by known spectral information. Calibration may be done using a nanohole array illuminated by a white light filtered by a set of narrow-width band-pass filters, which allows us to define the relationship between the centroid pair distance and optical wavelengths. In grating-based sSMLM, this distance-wavelength relationship is linear, whereas, in prism-based sSMLM, the relationship is described by an empirical equation [27]. Additional axial calibration is required in 3D sSMLM, including DWP-based sSMLM.

### Advantages, limitations, and applications of sSMLM

The primary advantage of sSMLM imaging is its highly multiplexed super-resolution imaging during a single image acquisition. Using sSMLM, researchers reported resolving four dyes with highly overlapping emission spectra simultaneously using a single excitation laser [1]. However, this comes at the expense of spatial precision compared with SMLM. Because each single-molecule emission must be split into spatial and spectral PSFs, the photon count of each spatial PSF is reduced, resulting in a lower spatial precision [7]. 4Pi sSMLM addressed this concern by using two objective lenses to effectively double the photon count [1]; however, it considerably complicated the optical alignment and required transparent, thin samples. While DWP-based sSMLM also splits photons for spatial and spectral imaging, it has advantages over other sSMLM implementations. DWP-based sSMLM is less lossy than the single-objective prism approach, which has multiple glass-air interfaces [3], and the single-objective diffraction grating approach, which loses photons to unused diffraction orders [2]. As DWP is a prefabricated single module designed to be inserted into the side port of the microscope body, it is also simpler to implement than other approaches. Because the DWP is a prefabricated module, the distance between the dispersive element and the camera is fixed by design at the distance that yields the ideal level of spectral dispersion [27]. For other sSMLM approaches, spectral dispersion must be measured, and the position of the dispersive element must be adjusted multiple times during the system setup. Finally, DWP-based sSMLM enables 3D imaging thanks to the built-in 3D biplane design [14, 28].

It is also possible to adapt DWP-based sSMLM to perform 2D imaging by altering the DWP such that the right prism is slightly displaced forward by 1.5 mm. This reduces the path length of the spectral channel. While the spectral channel still experiences a slightly longer path length than the spatial channel, this is compensated by it traveling through longer pathlength in glass as shown in Fig. S1.

## System description

### Illumination path

The illumination of an sSMLM system is identical to that of an SMLM setup, as previously described [5, 29]. Briefly, in the illumination path (Fig. 1a), a 642 nm laser (or a laser at other wavelengths) is aligned using two mirrors (M1 and M2) through a 10× beam expander (BE, consisting of a 50-mm lens and a 500-mm lens in series 550 nm apart). The expanded beam is adjusted to the height of the microscope body (Ti2, Nikon) backport using a periscope (assembled from two elliptical mirrors mounted on a right-angle kinematic mount.) The light is then focused onto the back focal plane of a 100× objective lens using a 500-mm focusing lens. The objective lens is also compatible with the Nikon Perfect Focus System. In our setup, the periscope and lens are all on a Thorlab cage system for easy alignment. The light is directed into the objective lens using a filter cube containing a 640±14 nm band-pass filter followed by a dichroic mirror to reflect the laser light into the objective lens (LF635/LP-C-000-NTE, Semrock).

**Figure 1.**
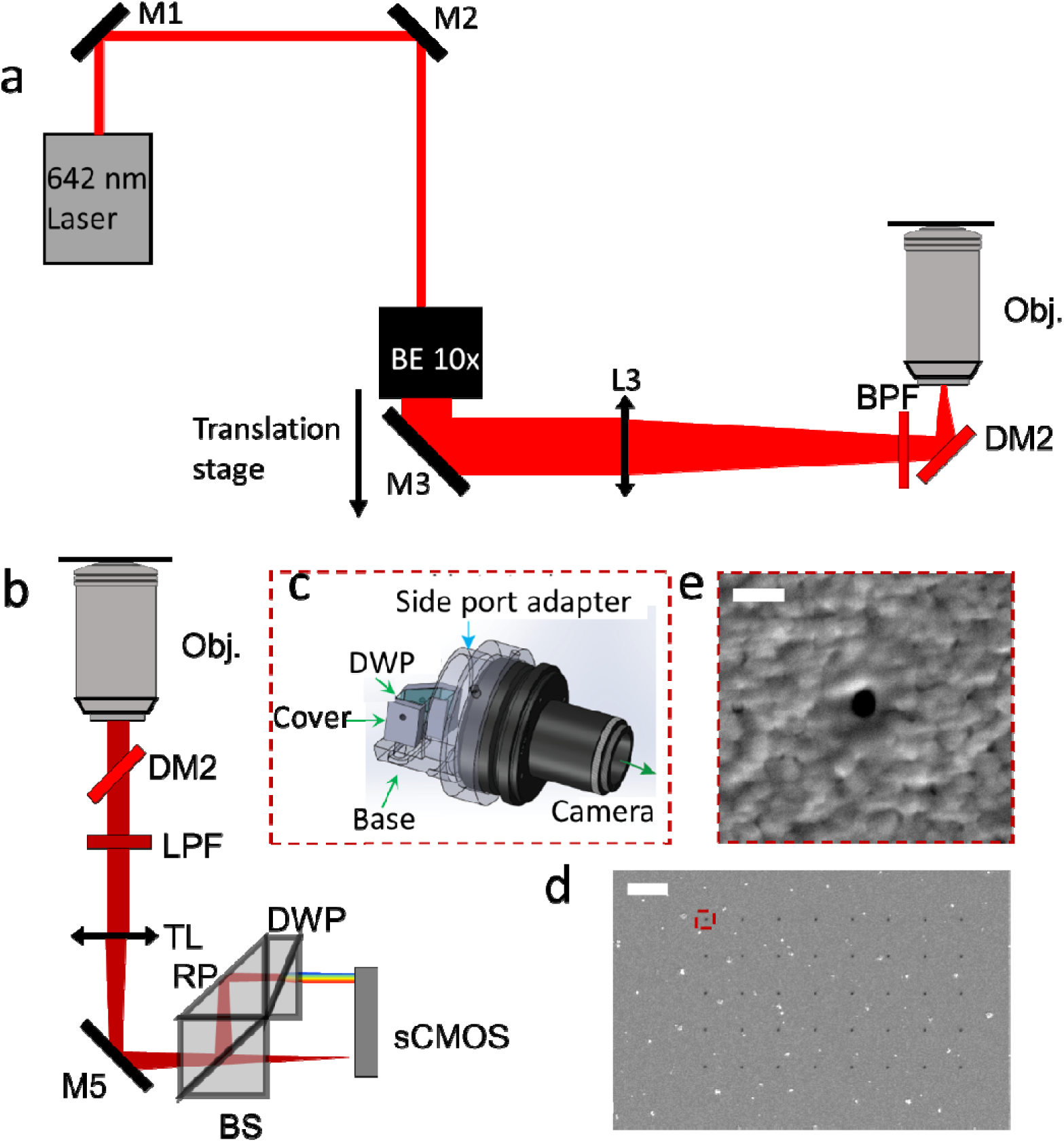
3D DWP sSMLM system detection and excitation path optical overview (drawn not to scale for clarity). (a) Adjustable illumination light path. M: mirror; DM: dichroic mirror; BE: Beam expander; L: lens, BPF: band-pass filter; Obj.: objective lens. (b) DWP sSMLM detection light path. LPF: long-pass filter; TL: tube lens; BS: Beam splitter; RP: right prism; DWP: dual-wedge prism. (c) Solidworks rendering of DWP assembly and its holder that slots into the sideport of the microscope body. (d) SEM image of the nanohole array, scale bar: 5 m. (e) Magnified view of one nanohole highlighted in panel (d), scale bar: 200 nm.

### Detection path

In the detection path (Fig. 1b), the fluorescence emissions are captured by the objective lens and passed through the previously mentioned dichroic mirror. They are focused by a tube lens inside the microscope body and then pass through a long-pass filter to remove residual excitation light. From this point on, the DWP-based sSMLM differs from the standard SMLM system, as the photons will go through the DWP assembly instead of being captured by an array detector. The DWP assembly consists of a nonpolarizing broadband beam splitter cube (BS), which splits the photons into a spatial path and a spectral path, a right prism (RP) that reflects the spectral image, and DWP that acts as the dispersive element. The spatial path forms an image on the camera, while the spectral path is reflected by a right prism into two wedged triangular prisms that collectively serve as the dispersive element. The DWP fits into a holder that slots into the microscope sideport (Fig. 1c) and can be connected directly to the sCMOS camera.

## Overview of key component fabrication and alignment procedure

### Fabrication of the DWP and DWP holder

Following our design, the DWP assembly was made by a third-party vendor (Hyperion Optics, Edison, NJ). The DWP itself consists of two cemented wedge prisms, referred to as WP1 (N-SF11 glass, refractive index n = 1.791 at 550 nm) and WP2 (N-LAF21 glass, refractive index n = 1.792 at 550 nm) [14]. The DWP assembly holder consists of three pieces (Fig. 1c), which are made from aluminum (6061-T6) and black anodized. The DWP assembly base, cover, and the circular adapter can slot into the microscope sideport.

### (Optional) Fabrication of the nanohole array (Steps 1-6)

Users can use a nanohole array to perform spectral calibration, where the array consists of a 5×8 rectangular lattice with a hole diameter of 100 nm and a spacing of 4 µm on an aluminum (Al)-coated glass coverslip. Fig. 1d is an SEM image of the nanohole array, and Fig. 1e shows a single hole highlighted by the red square in Fig. 1d. The nanohole array was fabricated using metal sputtering and focused ion beam (FIB) milling. First, a 150-nm-thick Al film was deposited on the borosilicate glass coverslip (22×22 mm^2^) by sputtering (ATC Orion Sputter System, AJA International Inc.) with a direct current (DC) power of 75 W and 20 standard cubic centimeter per minute (sccm) of argon (Ar) at 3 mTorr. Then, we used FIB milling (JIB-4700F FIB-SEM, JEOL Ltd.) with an acceleration voltage of 30 kV to fabricate the 5×8 nanohole array in the aluminum. Alternatively, performing spectral calibration using the United States Air Force (USAF) 1951 resolution chart (R3L1S4N, Thorlabs) instead of the nanohole array is possible. A sample image of the resolution chart used for the calibration procedure is shown in Fig. S2.

### Alignment of the excitation path (Steps 7-72)

We recommend mounting the entire sSMLM system on a floating optical table for mechanical stability (Fig. S3). First, install the microscope body and the desired lasers (steps 7-22). Use two mirrors (M1 and M2) to direct the lasers towards M3, then use M3 and M4 (Fig. S4) to direct the laser towards the location of the periscope system, leaving at least 550 mm between M4 and the periscope (Steps 23-26). Then place two irises (I1 and I2, as shown in Fig. S4) between M4 and the periscope location, screwing them into the breadboard holes to ensure that they are coaxial. Then, iteratively adjust M3 and M4 until the light passes through the center of both irises (Steps 27-36). Next, add a 10× beam expander, composed, in our case, of a 50-mm lens (L1) and a 500-mm lens (L2). Add the 50 mm lens on a base that is also screwed into the breadboard holes.

Then, adjust its height and rotation until the light passes through both irises again. Next, add the 500 mm lens and ensure collimation of the beam after the second lens by confirming that the beam size near the lens is equal to the beam size far from the lens (Steps 37-40). Next, add the periscope connected to a cage system (Fig. S5), all on a translational stage that moves left and right relative to the backport (Steps 41-50). Place a cage system iris (LCP50D, Thorlabs) in the cage system so that the center of the iris is aligned with the center of the microscope backport (Fig. S6). Next, move the iris between two set positions while iteratively adjusting the two periscope mirrors until the beam passes through the center of the iris at both positions (Steps 51-56). Next, insert the filter cube into the filter cube turret and rotate the turret to the backport position. Then, rotate the objective lens turret to a position with no objective lens and remove the protective cover. Iteratively adjust the periscope mirrors (EM1 and EM2 in Fig. S6) until the beam path passes through the center of the backport of the microscope and the center of the objective lens hole. Rotate the objective lens turret into the position so the beam passes through the objective lens. Finally, iteratively adjust the two periscope mirrors until the light passes through the center of the microscope backport and the exit of the objective lens (Steps 57-69). Place a 2-inch 500 mm focal length lens at the end of the cage system and adjust its position until the light coming out of the objective lens is as close as possible to collimation (Steps 70-72).

### Alignment of the detection path (Steps 73-96)

Install the camera and ensure the correct position by illuminating the nanohole array with white light and confirming the array size based on the objective lens magnification and camera pixel size (Steps 73-87). If no nanohole array is available, it is also acceptable to use a USAF 1951 resolution chart. Ensure that the sample is in focus when viewed through the camera, as well as when viewed through the microscope eyepiece. Similarly, ensure that the FOV is centered in the same place in the camera and in the microscope eyepiece. Assemble the filter wheel containing five band-pass filters, assemble the DWP, and insert them into their respective positions (steps 88-96).

### Spectral calibration (Steps 97-115)

We describe how to perform spectral calibration based on imaging a nanohole array illuminated by white light (the microscope lamp is acceptable). If no nanohole array is available, it is also acceptable to use a suitable line spacing of the USAF 1951 resolution chart mentioned above, and use the raw images in RainbowSTORM directly. Capture 100 frames through each position of the filter wheel, using a region-of-interest (ROI) containing the spatial (Fig. 2a) and spectral (Fig. 2b) images, and save them to an nd2 file. Localize the spatial and spectral images using the ThunderSTORM ImageJ plug-in [30, 31] to perform maximum likelihood estimation (MLE) Gaussian fitting. When setting up the ThunderSTORM parameters, use an intensity threshold of 3 times the standard deviation of the first wavelet level [32] for approximate localization. For sub-pixel localization, use a fitting radius of 330 nm and an initial sigma guess of 175 nm. Adjust the parameters as needed to ensure that only the desired nanoholes are localized. Save each localization list to a comma-separated value (csv) file in a subfolder. Next, using the RainbowSTORM v2 MATLAB app, click on Spectral Calibration, and a pop-up window named ‘Spectral Calibration’ should appear (Fig. 2c). Using this program, browse the subfolder where the csv files are saved, and the application automatically loads and processes the csv files, aligning the localizations between the zeroth and first orders and generating plots that display the shifts observed and expected at various wavelengths. If the wavelength detected for each file is incorrect, users can manually enter the correct wavelength. After these adjustments, save the calibration file by clicking the ’Save Calibration File’ button. This calibration file contains essential parameters, including the transformation data needed to correct any distortions between the zeroth and the first orders. This data also includes a calibration curve correlating the distance between spatial localizations and the corresponding spectral localizations The saved calibration file may be used for images obtained under identical imaging conditions.

**Fig. 2.**
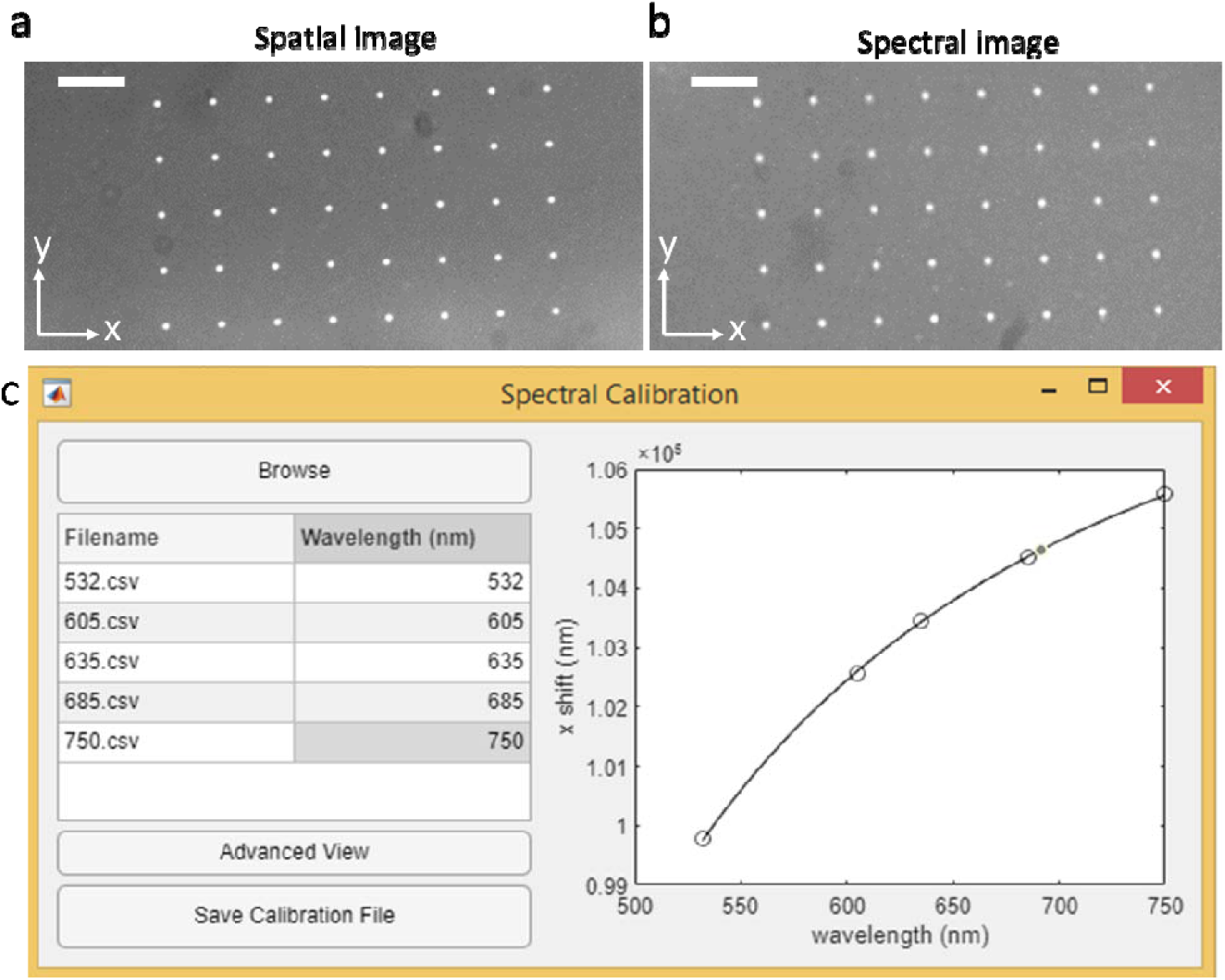
Spectral Calibration using a nanohole array. (a) The spatial image of the nanohole array and filtered by a 685 nm cutoff band-pass filter, scale bar: 5 m. (b) The spectral image of the nanohole array and filtered by a 685 nm cutoff band-pass filter, scale bar: 5 m. (c) The GUI of the Spectral Calibration menu in RainbowSTORM v2 with a graph showing a plot of distance between spectral and spatial localizations as a function of the cutoff wavelength of the band-pass filter.

### Axial calibration (steps 116-143)

We performed axial calibration by imaging microspheres with a mean diameter of 200 nm and an emission wavelength of 680 nm using the stack acquisition mode of Nikon’s NIS elements AR software and then the axial calibration feature in RainbowSTORM. Prepare a 1:100,000 dilution of microspheres in PBS and place 200 μL in a well of an 8-well chambered coverglass. Use Nikon NIS elements AR software for acquisition controls. First, use the widefield mode to illuminate the sample with 80 W/cm^2^ of 670 nm linearly polarized light for focus adjustment onto the desired ROI in the sample. Select a single nanosphere for calibration and ensure its PSFs are clearly visualized in the spatial and spectral images. Then, adjust the illumination brightness to ensure that the microsphere is bright (i.e., uses greater than 95% of the camera dynamic range) without saturating the sCMOS camera. Set the starting axial position such that the spatial image is slightly out of focus in the -z direction and the spectral image is slightly out of focus in the +z direction. Next, scan the microspheres across a 2 μm (±1 μm) axial range with an interval of 20 nm, ensuring that the PSF defocus is symmetric in the positive and negative z-direction. If the PSF defocusing is asymmetric, adjust the correction collar of the objective lens.

After imaging the microspeheres at all depths, open RainbowSTORM v2 and click on the “3D Processing” button and then the “Axial Calibration” button to open the Axial Calibration menu, from which users can specify the path to the nd2 file for axial calibration. The application will load the microsphere data and provide a preview of the spatial (Fig. 3a) and spectral (Fig. 3b) images. RainbowSTORM will sum the spatial image and spectral image along the x-direction, which should result in a roughly Gaussian-shaped 1D PSF for each channel (Figs. 3a&3b) since spectral dispersion is only in the x-direction. RainbowSTORM then uses MLE to fit a 1D Gaussian function to each channel and record the standard deviation σ at each axial position. At the end, RainbowSTORM generates the relationship between full-width-at-half-maximum (FWHM) values of the spatial and spectral PSFs and their corresponding z positions, where the separation between the minima of the two graphs corresponds to the difference in focal points (Fig. 3c). The correction collar in the objective lens can be adjusted to tune the separation to increase axial range at the expense of axial precision [7]. The cutoffs for the minimum (Z0) and maximum (Z1) axial positions may be adjusted using the sliders below the FWHM vs. z-position graph. From this graph, RainbowSTORM calculates the ratio between the spectral to spatial FWHM PSF sizes as a function of axial position (Fig. 3d). At the end, it saves the calibration file as a csv file, which contains the information needed to assign each localization a z-position in imaging data.

**Figure 3.**
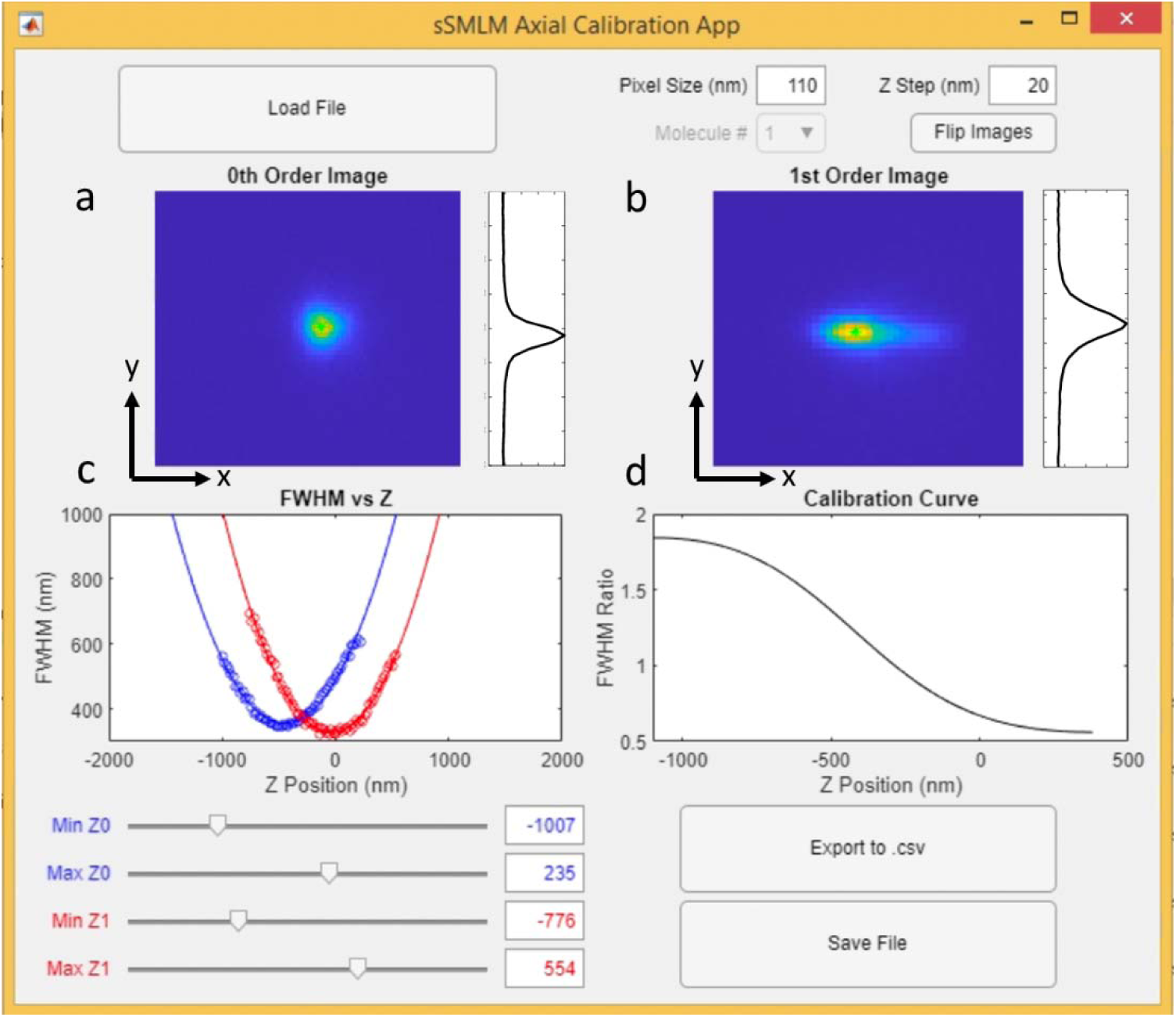
The RainbowSTORM v2 axial calibration window. (a) The spatial image of an isolated microsphere along with its sum in the *x*-direction. (b) The spectral image of an isolated microsphere along with its sum in the *x*-direction. (c) The FWHM of the spatial PSF (blue) and the spectral PSF (red). As a function of objective lens axial position, with sliders to adjust range . (d) The curve of spectral to spatial FWHM ratio as a function of objective lens axial position.

### Cell thawing and culturing (steps 144-176)

We use HeLa cells as the sample; users can use any cell type for specific investigations. First, thaw a vial of frozen cells in a warm water bath (37°C). Then, add the thawed contents to a separate tube and slowly add 4 mL of cell culture media (90% DMEM, 9% fetal bovine serum, and 0.9% penicillin-streptomycin). Centrifuge the cells at 400 g for five minutes, condensing them into a pellet. Finally, aspirate the supernatant, add 6 mL of cell culture media, and transfer the cells to a 25 cm^2^ cell culture flask. Store the cells at 37°C with 5% CO_2_ until they reach 60-80% confluency, then detach them from the flask by washing once with PBS, adding 1 mL trypsin-EDTA, and incubating the cells for five minutes at 37°C. Next, add 4 mL of cell culture media, centrifuge the cells at 400 g for five minutes, and condense them onto a pellet. Aspirate the supernatant, add 6 mL of cell culture media, and transfer 2 mL to a new flask for future use, supplementing with 3 mL of cell culture media. Transfer 200 μL of the cell suspension to each well of an 8-well chambered coverglass, wait until the cells reach 50-80% confluency (∼24 hrs), and proceed to cell fixation.

### Secondary antibody preparation (steps 177-192)

Reconstitute the selected fluorescence dye to a concentration of 0.2 mol/L in DMSO. Combine 1.2 μmol of dye, 0.13 mg of antibodies, and 10 μmol of NaHCO_3_ in a microcentrifuge tube, mix them well, and incubate at room temperature for 2 hours. Add 400 μL of PBS to an ultra-centrifugal unit and centrifuge at 14,000 g for 10 minutes. Move the ultra-centrifugal filter to another tube, transfer the contents of the microcentrifuge tube to the ultra-centrifugal unit, and centrifuge it at 14,000 g for 10 minutes. Add 200 μL of PBS to the ultra-centrifugal unit and centrifuge at 2,000 g for 5 minutes. The anti-rabbit AF647 was bought with the antibody already conjugated to the fluorophore and therefore the dyes and antibodies do not need to be conjugated.

### Cell fixation and labeling (steps 193-226)

After 24 hours, wash the cell-containing wells twice with PBS, then incubate them at room temperature in a fixative solution (3% paraformaldehyde and 0.1% glutaraldehyde in PBS) for 10 minutes. Wash the wells three times with PBS and incubate the cells in permeabilization buffer (0.5% triton X 100 in PBS) for 10 minutes at room temperature. Next, block the cells by incubating each well in blocking buffer (2.5% goat serum in PBS) for 30 minutes at room temperature. Perform primary antibody labeling using a primary antibody that targets the protein of interest. For multiplexed four-color imaging, incubate the cells in a solution composed of 1:300 mouse anti-pmp70 antibody, 1:200 rabbit anti-TOM20 antibody, 1:400 rat anti-vimentin antibody, and 1:200 chicken anti-tubulin antibody in blocking buffer overnight at 4°C. Then wash the cells three times with washing buffer (2% goat serum, 2% bovine serum albumin, and 0.1% triton X 100 in PBS) for 5 min each wash. Next, incubate the cells in the secondary antibody solution, in this case, a solution composed of 1:500 donkey anti-mouse DY634, 1:250 goat anti-rabbit AF647, 1:1500 donkey anti-chicken CF660, and 1:5000 donkey anti-chicken DyLight 680 (DL680) in blocking buffer for 45 minutes at room temperature. Wash twice with washing buffer for five minutes at room temperature and twice with PBS for five minutes at room temperature.

### Imaging (steps 227-276)

Prior to imaging, place the sample in imaging buffer (50 mM Tris HCl (pH=8.0), 10 mM NaCl, 0.5 mg/mL glucose oxidase, 2000 U/mL catalase, 10% (w/v) D-glucose, and 100 mM 2-Mercaptoethylamine Hydrochloride. For acquisition controls, use the Nikon NIS Elements AR software. First, use widefield to illuminate the sample using linearly polarized light of 642 nm at 80 W/cm^2^ to find and focus on the desired ROI in the sample. This is done because in preview mode, the structure of the sample is easier to see when the laser power is too low to induce blinking. A typical field-of-view (FOV) is 45×43 µm^2^. Then, ensure that the sample is fully visible in the spatial and spectral images and define the acquisition ROI to limit the size of the raw imaging data. After ensuring that the perfect focus is engaged, increase the illumination power to 6 kW/cm^2^ and acquire a video of the blinking fluorophores with a typical exposure time of 10 ms for 30,000 to 50,000 frames. The number of frames and the exposure time can be adjusted as needed. At the end of the acquisition, save the video as an nd2 file.

### Image analysis (steps 277-303)

Open the nd2 file in ImageJ. To reduce memory use, use Bio-Formats to only open the left 400 pixels when processing the spatial image. Likewise, when processing the spectral image, only open the right 400 pixels. Take note of the starting pixel values of the cropped image, and save each cropped image as a tiff. Use ThunderSTORM to perform localization in each tiff image based on MLE Gaussian fitting. For the spatial image, use an intensity threshold of 1.2 times the standard deviation of the first wavelet level [32] for approximate localization. For sub-pixel localization, use a fitting radius of 3 pixels and an initial sigma guess of 1.5 pixels. For localization of the spectral image, use an intensity threshold of 1.1 times the standard deviation of the first wavelet level, a fitting radius of 4 pixels, and an initial sigma of 2.5 pixels. The differences between fitting for the spatial and spectral images account for the fact that the spectral PSFs occupy more pixels and have a lower peak intensity than the spatial PSFs. Because spectral PSFs are larger than spatial PSFs, there is a tendency for a single spectrum to be recognized as two separate localizations. To avoid this, After localization of the spectral PSFs, use Thunderstorm to remove all localizations within 500 nm of each other within a single frame, ensuring only one localization for each spectral PSF. Export each list of localizations and statistics to csv files. Open RainbowSTORM and enter the file paths for the spectral and spatial csv files into their respective fields and the output path. If needed, enter the file path of the axial calibration file into its corresponding field. Adjust the sliding bars under the displayed image to set the spectral limits of each color channel [33].

### (Optional) Quantifying axial resolution

Users can quantify axial resolution by imaging 200 nm microspheres. Prepare a 1:100,000 dilution of microspheres in PBS and place 200 μL in a well of an 8-well chambered coverglass (12-565-470, Fisher Scientific). First, use the widefield mode to illuminate the sample with 80 W/cm^2^ of 670 nm linearly polarized light for focus adjustment onto the desired ROI in the sample. Select a single nanosphere for calibration and ensure its PSFs are clearly visualized in the spatial and spectral images. Adjust the illumination brightness as needed to ensure the microsphere emits between 300 and 4,000 photons per frame. At each axial position, adjust a continuous variable neutral density (ND) filter (NDC-50C-2M, Thorlabs) and the ambient light axial position in a 1 µm range at an incremental of 50 nm, acquiring 100 frames at each position, to perform multiple acquisitions at various illumination intensities. Then, manually adjust the and saving the frames to an nd2 file. Perform the previously described image analysis to the axial resolution data to extract the z-position in each frame. Next, compute the standard deviation of the estimated z-coordinate at each axial position, which yields the axial precision. To calculate axial resolution, multiply the axial precision by 2.355 [34]. Next, use least squares fitting to fit this to the equation for axial precision via biplane imaging [7]

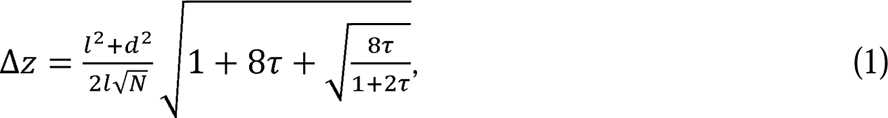

where 2*l*, [nm] is the mutual defocus of the emmitters in each channel; *d* is the focal depth; *N* [-] is the number of photons; and *τ* is defined as

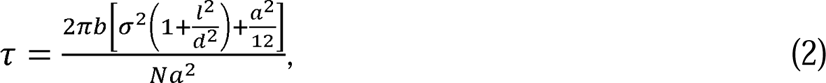

where *b* [-] is the number of background photons; *σ* is the standard deviation [nm] of the PSF; and *a* [nm] is the pixel size. Tune the parameter *d*, until the theoretically calculated axial precisions most closely match the measured ones. The equation with the tuned *d* may then be used to estime axial resolution in experimental datasets.

## Detailed description of the procedure

### Fabrication of the nanohole array (Optional)

1. Prepare the 22×22 mm^2^ borosilicate glass substrate.
2. Remove any organic materials from the glass substrate by informal cleaning with Nano-Strip®, a stabilized formulation of sulfuric acid and hydrogen peroxide compounds.
3. Rinse the glass substrate with deionized (DI) water 7 times and remove the remaining water from the substrate by nitrogen gas blowing.
4. Dehydration of the substrate on the hotplate at 110°C for 30 minutes.
5. Deposit an Al layer with a thickness of 150 nm on the borosilicate glass substrate by sputtering (ATC Orion Sputter System, AJA International Inc.). The detailed sputtering conditions are as follows: DC power is 75 W. Ar is 20 sccm. Pressure is 3 mTorr. The working distance between the Al target and the substrate is 35 mm. The deposition rate ∼ 1.5 nm/min.
6. Fabricate the nanohole array on the Al film on the glass substrate by FIB milling (JIB-4700F FIB-SEM, JEOL Ltd.). The nanohole array is designed to be a 5×8 rectangular lattice with a diameter of 100 nm and a period of 4 µm. The acceleration voltage is 30 kV.

### Excitation alignment procedure

7. Set up a floating optical table over 4 feet by 8 feet in a room without natural light.
8. Place the Ti2 microscope body on the floating table such that the eyepiece protrudes off the edge of the floating table and leaving sufficient room for the excitation and detection hardware (Fig. S3)
9. Ensure that the microscope is aligned with the floating table holes by checking that the front and back plates are not slanted compared to the breadboard holes, and fix it by screwing the front and black plates into the breadboard.
10. Set up a computer in the same room, ensuring the mouse is not resting on the optical table.
11. Install MPBC Laser control on the computer following the instructions provided by MPB Communications. Users can integrate other lasers into the system as well.
12. Set up the 642 nm laser following the instructions provided by MPB Communications.
13. Attach the laser module to four optical posts in 1.5” pedestal post holders using the four screw indentations in the laser module stand.
14. Place the 642 nm laser module perpendicular to the microscope body back port (Ls-642 in Fig. S4).
15. Fix the laser to the breadboard using clamping forks.
16. Turn on the laser by switching on the external power module.
17. Turn the key on the power module to the “on” position.
18. Wait until the MPBC Laser control indicates that the SHG temperature has reached its maximum point.
19. Choose current as the power control unit by clicking on the circle next to “Current, mA”.
20. Enter the minimum current possible that still allows the laser to emit light.
21. Click “Activate”.
22. Turn the 642 nm laser on and modulate its power to a safe level using a continuous variable ND filter attached to a 1” optical post in a 1.5” pedestal post holder.
23. Add M1 (Fig. S4) in a 1” kinematic mirror mount on a 1” optical post in a 1.5” pedestal post holder so that the laser light is roughly parallel to the microscope backport.
24. Add M2 (Fig. S4) in a 1” kinematic mirror mount on a 1” optical post in a 1.5” pedestal post holder so that the laser light is roughly parallel to its original orientation.
25. Add M3 (Fig. S4) in a 1” kinematic mirror mount on a 1” optical post in a 1.5” pedestal post holder so that the laser light is roughly parallel to the microscope backport.
26. Add M4 (Fig. S4) in a 1” kinematic mirror mount on a 1” optical post in a 1.5” pedestal post holder so that the laser light is roughly parallel to its original orientation.
27. Add an iris (I1) on a 1” optical post in a 1.5” post holder both screwed into the same row of holes on the floating table matching Fig. S4.
28. Adjust the I1 center height to roughly the height of the center of M4.
29. Adjust the knobs on the M3 kinematic mount until the laser light passes through the center of the iris.
30. Place and tighten a post collar on the I1 post.
31. Screw a 1” post holder into the optical table at the position indicated for I2 in Fig. S4.
32. Remove the I1 post and post-mounted iris and place it in the I2 post holder. The collar will ensure it is the same height.
33. Place the second iris on a 1” post in the I1 post-holder and adjust its height until the laser passes through the center of the iris.
34. Adjust the M4 mount knobs until the laser light passes through the center of I2
35. Adjust the M3 mount knobs until the laser light passes through the center of I1
36. Iteratively repeat steps 34-35 until the light passes through the center of I1 and I2
37. Add the 50 mm focal lens (L1, Fig. S4) on a 1” lens holder on a 1” post in a 1.5” pedestal post holder in front of I1 and in front of the beam. Adjust the height, angle, and position until the beam passes through the center of I1 and I2.
38. Screw L1 into the optical table.
39. Add a 500 mm focal lens (L2, Fig. S4) on a 2” lens holder on a 1” post on a 1.5” post holder before I2 and 550 mm after L1. Adjust the height, angle, and position until the beam passes through the center of I2. Adjust the distance between L1 and L2 until the beam is fully collimated after L2. Check using a piece of paper to verify collimation.
40. Fix L2 to the optical table using a clamping fork.
41. Screw a translational stage (TS) into the optical table along the same axis as the irises.
42. Screw an aluminum breadboard (BB) in on top of the TS (Fig. S5a).
43. Insert the elliptical mirrors (EM1 and EM2) into the elliptical mirror mounts and tighten the tightening screws.
44. Construct the periscope system out of 2 elliptical mirrors in mounts connected by 4 1.5” cage assembly rods (Fig. S5b).
45. Attach two upside-down 4” pedestal posts each to a horizontal cage clamp (CC).
46. Attach the 4” pedestal posts each to an upright pedestal post (Fig. S6).
47. Attach each 6” cage assembly rod to a 1” cage assembly rod.
48. Attach the assembly rod constructions to the periscope as in Fig. S6.
49. Attach the assembly rod-periscope to the horizontal cage clamp as in Fig. S6.
50. Clamp the pedestal posts to the breadboard using the large clamping forks as in Fig. S6.
51. Add the 60 mm cage iris to the upper cage system on the periscope.
52. Adjust the cage clamp height until the center of the iris matches the center of the microscope back port.
53. If the laser light does not hit roughly the center of the lower elliptical mirror of the periscope, adjust the height of all lenses, irises, and mirrors and repeat step 36.
54. Adjust the EM1 mount knobs until the laser light passes through the center of the 60 mm cage iris.
55. Adjust the EM2 mount knobs until the laser light enters the center of the microscope back port.
56. Iteratively repeat steps 47-48 until the laser light passes through the center of the 60 mm cage iris and the center of the microscope back port.
57. Construct a 642 nm laser filter cube out of the Nikon filter cube, the 640/14 nm band-pass filter facing the microscope back port, the 635 nm edge dichroic mirror across the center of the cube, and a 635 nm cutoff long pass filter facing the sample holder, as in Fig. S7a.
58. Slot the 642 nm laser filter cube in the microscope filter turret and turn it until the 640/14 nm band-pass filter faces the microscope back port.
59. Remove the objective lens cover from the microscope objective turret and rotate the turret until the hole is over the 642 nm laser filter cube.
60. Use a piece of tissue paper over the objective turret hole to test whether the laser light is coming out of the center of the hole.
61. Adjust the EM2 mount knobs until the laser light passes through the center of the objective turret hole.
62. Adjust EM1 mount knobs until the laser light passes through the center of the 60 mm cage iris.
63. Iteratively repeat steps 61-62 until the laser light passes through the center of the 60 mm cage iris at all positions along the cage and the center of the objective turret hole.
64. If the light is not passing through the center of the microscope back port, adjust the periscope height until it does.
65. Iteratively repeat steps 61-64 until the laser light passes through the center of the 60 mm cage iris at all positions along the cage, the center of the objective turret hole, and the center of the microscope back port.
66. Screw the 100× objective lens into the objective turret hole.
67. Adjust EM2 mount knobs until the laser light comes out of the objective lens with no angle.
68. Adjust EM1 mount knobs until the laser light passes through the center of the 60 mm cage iris.
69. Iteratively repeat steps 67-68 until the laser light passes through the center of the 60 mm cage iris and comes out of the objective lens with no angle.
70. Add a 500 mm focal lens (L3) on a 2-inch lens holder and place it on the end of the cage system close to the microscope back port.
71. Ensure that the laser light still comes out of the same place in the objective lens.
72. Move the lens towards and away from the back port until the beam size coming out of the objective lens is minimized.

### Detection path alignment procedure

73. Plug in and install the camera following instructions from photometrics.
74. Install Nikon NIS Elements AR/BR following instructions from Nikon.
75. Using the threaded holes in the camera, attach the camera to two 1” posts in 1” pedestal post holders.
76. Place the camera facing the desired microscope side port (Fig. S8) roughly 100 mm from the side port.
77. Turn on the microscope and the microscope white lamp.
78. Place a drop of immersion oil on the objective lens.
79. Place the nanohole array on the sample holder and with the center of the cross centered on the objective lens.
80. Adjust the focus until the immersion oil contacts the aluminum.
81. Switch the light so that it comes out of the eyepiece.
82. Adjust the x- and y-positions until the center of the cross is visible through the eyepiece.
83. Adjust the focus until the edges of the cross become sharp.
84. Adjust the x- and y-positions until a nanohole array is visible in the center of the eyepiece field of view.
85. Switch the light so that it comes out of the camera port.
86. Adjust the camera height and position until the nanohole array is visible on the left side of the camera field of view and center height.
87. Move the camera towards or away from the microscope body until the nanohole array appears as the correct size on the camera (the width of the nanohole array is 40 μm and the pixel size is 110 nm, so it should be 364 pixels wide) and the focus on the camera is the same as the focus in the eyepiece.
88. Use clamping forks to fix the camera position.
89. Place the DWP inside the DWP holder and tighten the positioning screw.
90. Slot the DWP holder inside the camera port, as in Fig. S8.
91. Assemble the filter wheel out of the filter wheel screwed onto a 1” post in a 1” pedestal post holder. Insert the following filters clockwise or anti-clockwise, ensuring that the arrow on each filter faces toward the camera:

a. Empty slot
b. 532 nm laser line filter
c. 580 nm bandpass filter
d. 633 nm laser line filter
e. 680 nm bandpass filter
f. 750 nm bandpass filter
92. Fix the filter wheel between the DWP and the camera using a clamping fork, as in Fig. S8.
93. Rotate the wheel so that the empty slot is facing the camera.
94. Turn on the camera, and open NIS elements. Press live view and set the LUT to autoscale.
95. Rotate the DWP holder until the spatial image is visible on the left side of the screen and the spectral image is visible on the right. Ensure that the spectra are straight across the x-axis with no vertical tilt by examining the pixels.
96. Tighten the screws in the camera port to fix the DWP holder position.

### Spectral calibration

97. Place the nanohole array target or the United States Air Force (USAF) Resolution Target on the sample holder of the microscope.
98. Rotate the filter such that the 532 nm laser line filter is facing the camera.
99. Select “Time acquisition” with 100 frames and an exposure time of 100 ms.
100. Enter the desired file name and file path.
101. Select “Run now”.
102. Repeat steps 98-101 with each band-pass and laser line filter in the filter wheel.
103. If using the USAF resolution target for spectral calibration, skip to Step 114.
104. Open Fiji and on the main application window, select the “plug-ins” dropdown list, the “BioFormats” dropdown sublist, and click on “BioFormats Importer”.
105. Choose the desired nd2 file containing the image filtered through the 532 nm band-pass filter.
106. Use the “Rectangle tool” to draw a square around the spatial and spectral image of the nanohole array.
107. On the main application window, select the “Plug-ins” dropdown list, the “ThunderSTORM” dropdown sublist, and click on “Run analysis”.
108. In the run analysis window, select “Camera setup” and ensure that the camera parameters match the specifications of the camera used for imaging.
109. Set the filter mode to Wavelet filtering with a B-spline order of 3 and a B-spline scale of 2.0.
110. Set the approximate localization method as Local maximum, set the peak intensity threshold as “3*std(Wave.F1), and set the connectivity to 8-neighborhood.
111. Set the sub-pixel localization method to “PSF: Gaussian”, set the fitting radius to 330 nm divided by the camera pixel size, set the fitting method to “maximum likelihood”, and set the initial sigma to 165 nm divided by pixel size. Run the analysis.
112. Click “Export results”, and save the results to a csv file.
113. Repeat steps 103-112 for each spectral calibration image.
114. Download the MATLAB app RainbowSTORM v2 from https://github.com/FOIL-NU/sSMLM-protocol.
115. Open the app and click on the “Spectral Calibration” button under the “Preliminaries” header.
116. In the Spectral Calibration menu, press the browse button and navigate to the folder where the spectral calibration csv files are saved.
117. After the data loads, adjust the wavelengths assigned to each csv file as needed, then click the “Save Calibration File” button.

### Axial calibration

118. Place 200 μL of PBS in four 1.5 mL microcentrifuge tubes labeled 1-4.
119. Place 1 μL of 200 nm 680 nm microspheres in microcentrifuge tube 1 and mix the solution.
120. Remove 1 μL of solution from microcentrifuge tube 1 and place it in microcentrifuge tube 2 and mix the solution.
121. Continue the serial dilution for tubes 3 and 4.
122. Transfer the solutions from each microcentrifuge to corresponding wells in an 8-well chambered coverglass.
123. Incubate the 8-well chambered coverglass for 30 minutes at room temperature.
124. Remove the solution from each well.
125. Add 200 μL PBS in each well.
126. Place a drop of immersion oil on the objective lens.
127. Place the 8-well chambered coverglass on the sample holder.
128. Rotate the filter wheel until the empty slot is between the DWP holder and the camera.
129. Adjust the *x*-*y* positions until the objective lens is under the well with solution from microcentrifuge tube 1.
130. Switch the light so that it comes out of the camera port.
131. Turn on the camera, and open NIS elements. Press live view and set the LUT to autoscale.
132. Turn off the white lamp and turn on the 642 nm laser (Steps 10-16).
133. Adjust the focus until the image is clear.
134. Adjust the *x*- and *y*-positions until the objective lens is under a well with highly sparse microspheres, adding more oil when necessary.
135. Select “Z acquisition” and select “symmetric mode defined by range”. Set a range of 2 μm and an exposure time of 10 ms. Set the center position to “relative” to ensure that the center *z*-position matches the current z-position displayed on the microscope.
136. Enter the desired file name and file path.
137. Adjust the *x*- and *y*-positions until at only one well-defined and sparse microsphere is present in the spatial and spectral images.
138. Adjust the *z* position to be between the focus of the spatial and spectral PSFs.
139. Ensure that the microsphere uses a significant portion of the camera dynamic range but does not saturate the camera. Adjust laser brightness on the laser control software or by adjusting the ND filter if necessary.
140. Select “Run now”.
141. Open the RainbowSTORM v2 app and click on the “3D Processing” button under the “Preliminaries” header.
142. Click on the “Axial Calibration” button under the “Preliminaries” header.
143. In the Axial Calibration menu, press the “Load file” button and navigate to the folder where the axial calibration nd2 or tiff file is saved and open it.
144. Adjust the “Min z0”, “Max z0”, “Min z1”, and “Max z1” sliding bars until the “FWHM vs Z” graph shows two clear parabolic curves.
145. Press the “Export to CSV” button.

### Cell thawing and culturing

146. Prepare a balance 15 mL centrifuge tube with 6.5 mL of water or PBS.
147. Screw the filter onto the 500 mL bottle that comes with it, and attach the filter nozzle to the cell culture hood vacuum and turn on the vacuum.
148. Add 500 mL DMEM, 50 mL fetal bovine serum, and 5 mL pen-strep in succession to the filter to make cell culture media.
149. Remove the filter from the 500 mL bottle and replace it with the bottle cap.
150. Remove two frozen cell vials from dry ice and place it in a latex glove. Tie off the opening.
151. Place the latex glove in the warm water bath set to 37 °C, until the cells are thawed.
152. Inside the cell culture hood, use the pipet aid with a 5 mL pipette to remove the cells from the frozen cell vials and place them in a 15 mL centrifuge tube.
153. Use the pipette aid with a 5 mL pipette to add cell culture media drop by drop to the cells in the 15 mL centrifuge tubes.
154. Place the balance and the 15 mL centrifuge tube containing cells on opposite sides of the tabletop centrifuge.
155. Spin the cells for 5 minutes at 800 g.
156. Remove the cell centrifuge tube from the centrifuge and place it in the cell culture hood.
157. Aspirate the supernatant from the 15 mL centrifuge tube using a glass Pasteur pipette connected to the cell culture hood vacuum, leaving the cell pellet.
158. Use the pipette aid with a 10 mL pipette to add 6 mL of cell culture media to the 15 mL centrifuge tube, homogenizing the cell pellet in the solution.
159. Transfer the solution to a cell culture flask.
160. Store the cell culture flask in the incubator at 37 °C, with 5% CO_2_.
161. view the cells daily with the tabletop light microscope until they reach 60-80% confluency.
162. Place the cells in the cell culture hood and aspirate the contents using a glass Pasteur pipette connected to the cell culture hood vacuum.
163. Use the pipette aid with a 5 mL pipette to add 1 mL of PBS to the cell culture flask.
164. Rock the flask to coat the bottom with PBS.
165. aspirate the contents of the cell culture flask using a glass Pasteur pipette connected to the cell culture hood vacuum.
166. Use the pipette aid with a 5 mL pipette to add 1 mL of trypsin-EDTA to the cell culture flask.
167. Rock the flask to coat the bottom with trypsin-EDTA.
168. Store the cell culture flask in the incubator at 37 °C, with 5% CO_2_ for five minutes.
169. view the cells daily with the tabletop light microscope to ensure the cells are detached from the cell culture flask; if they are not, place them in the incubator for another minute.
170. Place the cell culture flask inside the cell culture hood.
171. Use the pipette aid with a 5 mL pipette to add 4 mL of cell culture media to the cell culture flask.
172. Use the pipette aid with a 5 mL pipette to add 2 mL of solution from the cell culture flask to a new, empty cell culture flask.
173. Use the pipette aid with a 5 mL pipette to add 4.6 mL of cell culture media from the cell culture flask to the new cell culture flask.
174. Transfer 200 µL of solution from the new cell culture flask to each well of an 8-well chambered coverglass.
175. Discard the old cell culture flask and its contents using the appropriate waste management procedures.
176. Place the 8-well chambered coverglass and the new cell culture flask (for future use) in the incubator at 37 °C, with 5% CO_2_.
177. view the cells daily with the tabletop light microscope until they reach 60-80% confluency.
178. Proceed to cell fixation with the 8-well chambered coverglass.

### Secondary antibody preparation

179. Reconstitute each dye to a concentration of 0.2 mol/L in DMSO.
180. Place a 50 mL centrifuge tube on the scale and zero it.
181. Use the scale to weigh out 4.2 g of NaHCO_3_ in the tube.
182. Use the pipette aid with a 50 mL pipette to place 50 mL of PBS in the centrifuge tube and mix well to create 1M NaHCO_3_ solution.
183. Place 40 μL of 1M potassium hydroxide in the microcentrifuge tube. Add 100 μL of the antibody to a microcentrifuge tube.
184. Add 6 μL of the reconstituted dye to the microcentrifuge tube.
185. Add 10 μL of the 1 M NaHCO_3_ solution to the microcentrifuge tube.
186. Mix the microcentrifuge tube well.
187. Incubate the microcentrifuge tube at room temperature for 2 hours on the cell rocker.
188. Assemble the upper and lower chamber of the ultra centrifugal unit.
189. Add 400 μL PBS to the ultra filter (the upper chamber of the ultra centrifugal unit) and centrifuge at 14,000 g for 10 minutes.
190. Move the ultra centrifugal filter to another tube and transfer the contents of the microcentrifuge tube to the ultra centrifugal filter.
191. Centrifuge the ultra centrifugal unit at 14,000 g for 10 minutes.
192. Add 200 μL PBS to the ultra centrifugal filter and centrifuge at 2,000 g for 5 minutes.
193. Remove the filter and close the lid of the lower chamber.
194. (Optional) measure the absorption peaks of the antibody-dye conjugate to ensure roughly the desired ratio (2:1) of dye to antibody.

### Prepare fixation buffer

195. Use the pipette aid with a 10 mL pipette to add 6 mL 4% PFA to a 15 mL centrifuge tube.
196. Use the pipette aid with a 5 mL pipette to add 2 mL PBS to the 15 mL centrifuge tube.
197. Add 32 µL of 25% Glutaraldehyde to the 15 mL centrifuge tube (scale the amount of reagents up or down as needed).

### Prepare permeabilization buffer

198. Prepare a 10% triton X 100 stock buffer by adding 5 mL triton X 100 and 45 mL PBS to a 50 mL centrifuge tube.
199. Pour 14.25 mL of PBS in a 15 mL centrifuge tube.
200. Add 750 µL of the 10% triton X 100 stock buffer to the 15 mL tube.

### Prepare washing buffer

201. Prepare a 10% goat serum stock solution by adding 5 mL goat serum and 45 mL PBS to a 50 mL centrifuge tube.
202. Prepare a 10% bovine serum albumin stock solution by adding 5 mL bovine serum albumin and 45 mL PBS to a 50 mL centrifuge tube.
203. Add 8.85 mL PBS to a 15 mL centrifuge tube.
204. Add 3 mL 10% goat serum stock solution to the 15 mL centrifuge tube.
205. Add 3 mL 10% bovine serum albumin stock solution to the 15 mL centrifuge tube.
206. Add 150 µL of the 10% triton X 100 stock solution to the 15 mL centrifuge tube.

### Prepare blocking buffer

207. Add 300 µL PBS to a 1.5 mL microcentrifuge tube.
208. Add 100 µL 10% goat serum stock solution to the microcentrifuge tube.

### Cell fixation and immunolabeling

209. Remove the cell culture media from the well of interest and wash twice with PBS.
210. Add 200 µL of fixation buffer to the well of interest.
211. Incubate the cells at room temperature for 10 minutes.
212. Wsh the cells three times with PBS.
213. Remove the PBS and a add 200 µL permeabilization buffer to the well of interest and incubate for 10 minutes at room temperature on the cell rocker.
214. Remove the permeabilization buffer and add 200 µL permeabilization buffer to the well of interest and incubate for 10 minutes at room temperature.
215. Add 200 µL of blocking buffer to the well of interest and incubate for 30 minutes at room temperature on the cell rocker.
216. While waiting, add the primary antibody to the remaining blocking buffer. For example:

a. If labeling TOM20, use the 10 µL micropipette with a 10 µL micropipette tip to add 2 µL rabbit anti-TOM20 antibody.
b. If labeling histones, use the 10 µL micropipette with a 10 µL micropipette tip to add 1 µL rabbit Tri-Methyl-Histone H3 (Lys27) antibody and 1 µL mouse Histone H3K27ac Antibody.
217. Remove the blocking buffer from the well of interest and add the remaining blocking buffer with primary antibodies.
218. Place the cells on the cell rocker and incubate overnight at 4°C.
219. Remove all solution from the well of interest and add add 200 µL washing buffer and incubate on the cell rocker for 5 minutes at room temperature.
220. Repeat step 219 two more times.
221. While waiting:

a. Add 150 µL PBS to a 1.5 mL microcentrifuge tube.
b. Add 50 µL 10% goat serum stock solution to the microcentrifuge tube.
222. Add the secondary antibody to the remaining blocking buffer. For example

a. If labeling TOM20, use the 10 µL micropipette with a 10 µL micropipette tip to add 1 µL goat anti-rabbit AF647.
223. Place on the cell rocker and incubate for 45 minutes at room temperature.
224. Remove all solution from the well of interest and add 200 µL washing buffer and incubate on the cell rocker for 5 minutes at room temperature.
225. Repeat step 224.
226. Remove all solution from the well of interest and add 200 µL PBS and incubate on the cell rocker for 5 minutes at room temperature.
227. Repeat step 226.
228. Incubate the sample in the fridge with PBS in the well of interest until it is ready to be imaged.

### Preparing Tris buffer

229. Place a 1000 mL glass bottle on the scale and zero it.
230. Use the scale to weigh out 292.2 mg of NaCl and add it to the glass bottle.
231. Pour 475 mL of water into the bottle.
232. Add 15 mL of 1M Tris HCl to the glass bottle to make a 50 mM Tris HCl and 10 mM NaCl solution.

### Preparing 1 M potassium hydroxide solution

233. Place an empty 50 mL centrifuge tube on the scale and zero it.
234. Use the scale to weigh out 2.81 g of KOH in the tube.
235. Fill the tube to 50 mL with Tris buffer and mix it well.

### Preparing MEA buffer

236. Place a 1.5 mL microcentrifuge tube on the scale and zero it.
237. Use the scale to weigh out 113 mg of 2-Mercaptoethylamine Hydrochloride in the tube.
238. Place 1000 µL of tris buffer in the microcentrifuge tube and mix well.
239. Place 40 µL of 1M potassium hydroxide in the microcentrifuge tube.
240. Store the MEA buffer up to three weeks at 4 °C.

### Preparing glucose oxidase buffer

241. Place a 1.5 mL microcentrifuge tube on the scale and zero it.
242. Use the scale to weigh out 25 mg of glucose oxidase in the tube.
243. Place 1000 µL of tris buffer in the microcentrifuge tube and mix well.
244. Store the glucose oxidase buffer at 4°C.

### Preparing Tris + glucose buffer

245. Place an empty 50 mL centrifuge tube on the scale and zero it.
246. Add 5 g D-glucose to the tube.
247. Pour 45 mL tris buffer into the tube and shake until the glucose is dissolved.
248. Store the Tris + glucose buffer at room temperature.

### Preparing catalase buffer

249. Remove the metal cover from the lid of the catalase buffer.
250. Insert an 18-gauge 1.5” syringe and needle into the rubber cover.
251. Remove ∼0.5 mL of solution.
252. Transfer the solution into a 1.5 mL microcentrifuge tube.
253. Wait until the solution settles, forming a brown precipitate and a yellow supernatant.
254. Store the catalase buffer at 4°C.

### Preparing STORM buffer (prior to each imaging session)

255. Place 225 µL of Tris buffer in a 1.5 mL microcentrifuge tube and mix well.
256. Place 50 µL of MEA buffer microcentrifuge tube.
257. Place 5 µL of glucose oxidase buffer in the microcentrifuge tube.
258. Place 0.5 µL of catalase supernatant in the microcentrifuge tube.
259. Perform imaging within 3 hours. If more time is needed, prepare another batch of STORM buffer to replace the old buffer.

### Image acquisition

260. Prepare STORM buffer (see preparing STORM buffer section).
261. Remove all PBS from the well of interest in the 8-well chambered coverglass (See fixation and labeling section) and add 200 µL of STORM buffer to the well.
262. Turn on the microscope.
263. Place a drop of immersion oil on the objective lens.
264. Place the 8-well chambered coverglass on the sample holder.
265. Rotate the filter wheel until the empty slot is between the DWP holder and the camera.
266. Adjust the x-y position until the objective lens is under the well of interest.
267. Switch the light so that it comes out of the camera port.
268. Turn on the camera, and open NIS elements. Press live view and set the LUT to autoscale.
269. Turn on the 642 nm laser (Steps 10-16).
270. Adjust the focus until the fluorescent image is clear.
271. Adjust the x-y position until a desirable FOV appears.
272. Select “Time acquisition” with 30,000 frames and an exposure time of 10 ms.
273. Enter the desired file name and file path.
274. Ensure that the PFS is engaged.
275. In NIS elements, Draw an ROI containing the spatial and spectral image desired.
276. Select “Crop to ROI”.
277. Remove the ND filter and raise the current until the sample fluorophores appear to be blinking rapidly with sparse PSFs (it may be necessary to wait for the blinking density to go down).
278. Select “Run now”.

### Image analysis

279. Open Fiji and on the main application window, select the “Plug-ins” dropdown list, the “BioFormats” dropdown sublist, and click on “BioFormats Importer”.
280. Choose the desired nd2 file.
281. Under “Memory management”, select “Crop on import” then press “Ok”.
282. Set “x coordinate 1” and “Width 1” to align with the borders of the spatial image.
283. On the main application window, select the “plug-ins” dropdown list, the “ThunderSTORM” dropdown sublist, and click on “Run Analysis”.
284. In the run analysis window, select “Camera setup” and ensure that the camera parameters match the specifications of the camera used for imaging.
285. Set the filter mode to Wavelet filtering with a B-spline order of 3 and a B-spline scale of 2.0.
286. Set the approximate localization method as Local maximum, set the peak intensity threshold as “1.2*std(Wave.F1)”, and set the connectivity to “8-neighborhood”.
287. Set the sub-pixel localization method to “PSF: Gaussian”, set the fitting radius to 330 nm divided by the camera pixel size, set the fitting method to “maximum likelihood”, and set the initial sigma to 165 nm divided by pixel size. Run the analysis.
288. Click “export results”, and save the results to a csv file with the same name as the spatial image tiff file.
289. Repeat steps 279-281.
290. Set “x coordinate 1” and “Width 1” to align with the borders of the spectral image.
291. Repeat steps 283-286.
292. Set the approximate localization method as Local maximum, set the peak intensity threshold as “1.1*std(Wave.F1)”, and set the connectivity to “8-neighborhood”.
293. Set the sub-pixel localization method to “PSF: elliptical Gaussian (3D astigmatism)” using the calibration curve provided in the github, set the fitting radius to 440 nm divided by the camera pixel size, set the fitting method to “maximum likelihood”, and set the initial sigma to 275 divided by pixel size. Run the analysis.
294. On the results window, select “Remove duplicates”, set a distance threshold of 500 nm, and apply.
295. Click “export results”, and save the results to a csv file with the same name as the spectral image tiff file.
296. Open the RainbowSTORM v2 MATLAB app.
297. Under the “Directories” header, click on the “Browse” button next to the text that reads “0th Order Input File” and select the spatial image csv file ThunderSTORM output.
298. Under the “Directories” header, click on the “Browse” button next to the text that reads “1st Order Input File” and select the spectral image csv file ThunderSTORM output.
299. Under the “Directories” header, click on the “Browse” button next to the text that reads “Spectral Calibration File” and select the mat file generated from the spectral calibration procedure.
300. If doing 3D image processing, press the “3D Processing” button under the “Preliminaries” header.
301. Under the “Directories” header, click on the “Browse” button next to the text that reads “Axial Calibration File” and select the csv file generated from the axial calibration procedure.
302. Under the “Directories” header, click on the “Browse” button next to the text that reads “Output File” and select the desired name and file location of the output csv file.
303. Under the “Settings” header, shift the slider to the expected central wavelength of the emission wavelength of the fluorophores used in the sample.
304. Press the “Run!” button.

### Anticipated Results

Figure 4 shows typical results from 3D DWP-based sSMLM imaging. Figure 4a is a 2D projection of a four-color image [35] acquired with 3D DWP-based sSMLM of a HeLa cell using four red dyes excited simultaneously by a single laser, and Fig. 4b shows the histogram of estimated spectral peaks of the four dyes. With careful spatial and spectral calibrations, there is low cross-talk among the four colors (Figs. 4c-4f). Figures 4g & 4h show Figs. 4c & 4e with a color map corresponding to the axial position, indicating the axial distribution of the mitochondrial and tubulin networks [36].

**Figure 4.**
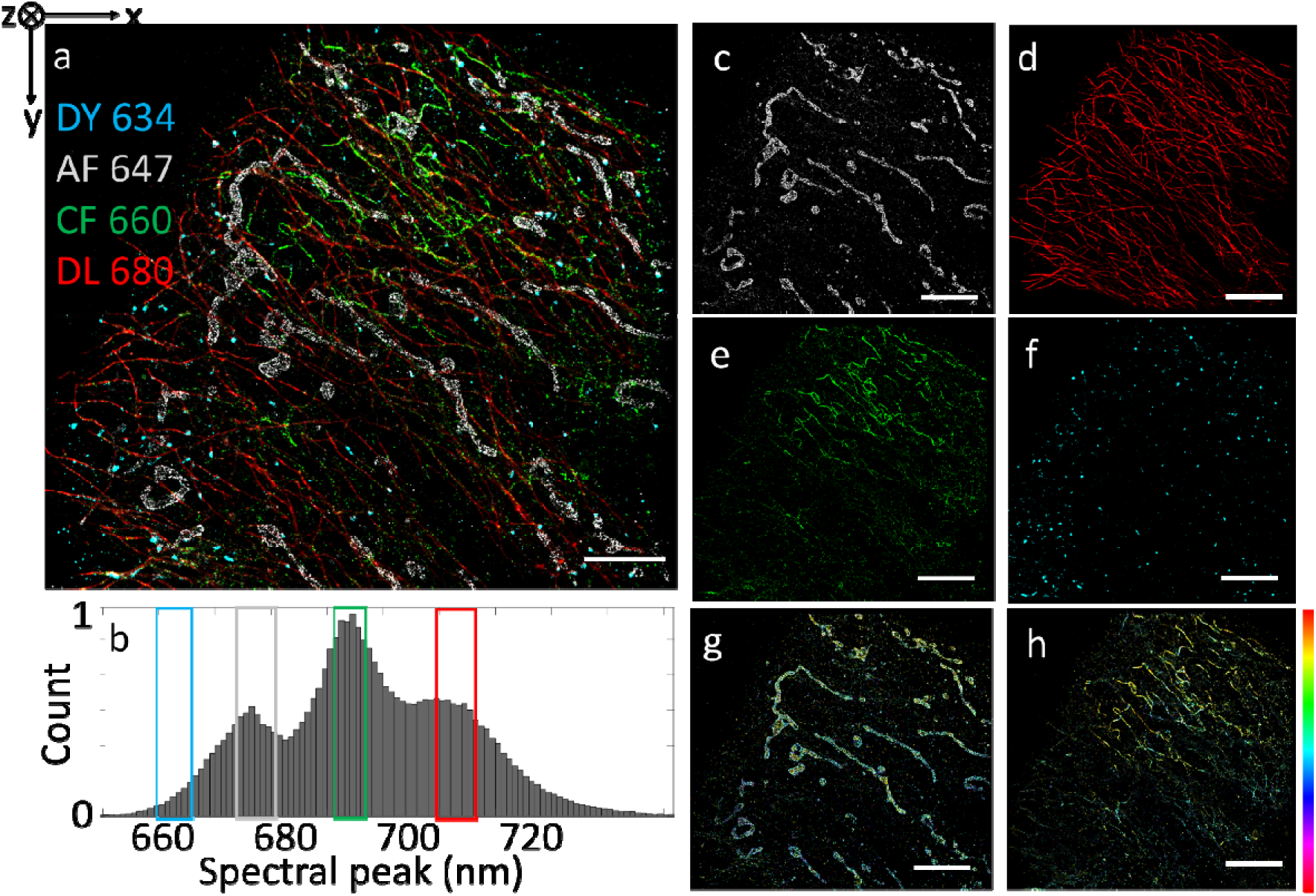
Typical imaging results. (a) 3D DWP 4-color image of a Hela cell immunolabeled using antibodies to alpha tubulin (DL680, red), vimentin (CF 660, green), Tom20 (AF 647, gray), and PMP70 (DY 634, cyan), scale bar: 5 μm. (b) A normalized histogram of spectral peaks identified using the spectral fitting and matching method. (c) Isolated mitochondria from (a) filtered by the spectral region shown by the grey square in (b) (scale bar: 5 μm). (d) Isolated tubulin from (a) filtered by the spectral region shown by the red square in (b). (e) Isolated vimentin from (a) filtered by the spectral region shown by the green square in (b). (f) Isolated PMP70 from (a) filtered by the spectral region shown by the cyan square in (b). (g) Same image as in panel (c) with pseudocolors corresponding to the axial position. Colorbar: -350 to 350 nm (h) Same image as in panel (e) with pseudocolors corresponding to the axial position.

## Materials

### Optomechanical equipment

- 18x clamping forks (CF125C, Thorlabs)
- 2x large clamping forks (PF175B, Thorlabs)
- 16x 1.5” pedestal post holders (PH1E, Thorlabs)
- 19x 1” posts (TR1, Thorlabs)
- 4x 1.5” post holders (PH1, Thorlabs)
- 4x 4” pedestal post (RS4P, Thorlabs)
- 1x 1” lens holder (CP33, Thorlabs)
- 2x 2” lens holder (LCP34, Thorlabs)
- 2x post-mounted irises (ID25, Thorlabs)
- 1x post collar (R2T, Thorlabs)
- 2x right angle kinematic elliptical mirror mounts (KCB2EC, Thorlabs)
- 4x 6” cage assembly rods (ER6, Thorlabs)
- 4x 1” cage assembly rods (ER1, Thorlabs)
- 4x 1.5” cage assembly rods (ER1.5, Thorlabs)
- 2x 60 mm cage horizontal clamp (CH1060, Thorlabs)
- 1x 2” translation stage (DTS50, Thorlabs)
- 1x 4” x 6” aluminum breadboard (MB4, Thorlabs)
- 5x 1” adjustable mirror mounts (POLARIS-K1E, Thorlabs)
- 1x Kinematic mount for rectangular optics (KM100C, Thorlabs)
- 1x filter wheel (KFW6, Thorlabs)
- 2x fluorescent filter cube (Nikon)
- 1x 60 mm cage system iris (LCP50D, Thorlabs)
- 1x 4’x8’x12.2” Optical table (T48WK, Thorlabs) Optical components
- 5x broadband dielectric mirrors (BB1-E02, Thorlabs)
- 2x 2” broadband dielectric elliptical mirrors (BBE2-E02, Thorlabs)
- 1x 56f1 nm edge dichroic mirror (Di02-R561-25x36, Semrock)
- 1x 561 nm cutoff long pass filter (BLP02-561R-25, Semrock)
- 1x 561/14 nm bandpass filter (FF01-561/14-25, Semrock)
- 1x 640/14 nm bandpass filter (FF01-640/14-25, Semrock)
- 1x 635 nm edge dichroic mirror (Di02-R635-25x36, Semrock)
- 1x 635 nm cutoff long pass filter (BLP01-635R-25, Semrock)
- 1x 532 nm laser line filter (FL532-3, Thorlabs)
- 1x 580 nm bandpass filter (FB580-10, Thorlabs)
- 1x 633 nm laser line filter (FL632.8-3, Thorlabs)
- 1x 680 nm bandpass filter (FB680-10, Thorlabs)
- 1x 750 nm bandpass filter (FB750-10, Thorlabs)
- 1x 1” 50 mm lens (47-637, Edmund optics)
- 2x 2” 500 mm lens (49-290, Edmund optics)
- 1x 100x objective lens (CFI Apochromat TIRF 100XC Oil, Nikon)

### Optical and computational equipment

- 1x Microscope body (Ti2, Nikon)
- 1x continuous wave 2W 642 nm laser (MPBC Communications)
- 1x sCMOS camera (Prime 95B, Photometrics).
- 1x 64-bit computer
- 1x Power meter (PM121D, Thorlabs)
- Immersion oil (Type F, Nikon) Software
- NIS Elements
- MPBC Laser control
- ImageJ (Fiji)
- ThunderSTORM
- RainbowSTORM v2
- MATLAB

### Cell Culture equipment

- 1x 8-well chambered coverglass (12-565-470, Fisher Scientific)
- 1x cell culture flask (156800, Thermofisher)
- 1x tabletop micro centrifuge (75004210, Thermofisher)
- 1x fully equipped cell culture hood (1300 series a2 thermo scientific)
- 1x 1000 µL micro-pipette (3123000063, Eppendorf)
- 1x 200 µL micro-pipette (3123000055, Eppendorf)
- 1x 10 µL micro-pipette (3123000020, Eppendorf)
- 1000 µL micro-pipette tips (13-811-159, Fisher Scientific)
- 200 µL micro-pipette tips (13-811-129, Fisher Scientific)
- 10 µL micro-pipette tips (13-811-105, Fisher Scientific)
- 1.5 mL microcentrifuge tubes (3451, Thermofisher)
- 5 mL pipettes (170355N, Thermofisher)
- 10 mL pipettes (170356N, Thermofisher)
- 15 mL centrifuge tubes (14-959-53A, Fisher Scientific)
- 50 mL centrifuge tubes (14-432-22, Fisher Scientific)
- 18-gauge 1.5” needle (AH+1838-1M, Nipro)
- 10 mL syringe (JD+10L2025-WEI, Nipro)
- Glass Pasteur pipettes (CLS9186, Sigma Aldrich)
- 1x Portable pipet aid (4-000-101, Drummond)
- 1x vacuum driven filter (5660020, ThermoFisher)
- 1x Water bath (15-462-10Q, Fisher Scientific)
- 1x incubator (11822711 Fisher Scientific)
- 1x Tabletop light microscope (invertoskop 40C, Zeiss)
- 1x Scale (Secura613-1s, Sartorius)
- 1x Cell rocker (benchmark mini BlotBoy, Tritech)
- 1 L Glass bottle (CH0164D, Eisco scientific)
- Ultra centrifugal filter (UFC501096, Sigma Aldrich) Reagents
- DMEM (11966025, Thermofisher)
- fetal bovine serum (SH3008802HI, Fisher Scientific)
- penicillin-streptomycin (15140122, Thermofisher)
- 2x tube of frozen HeLa cells (HTB-96, ATCC)
- trypsin-EDTA (T4174, Millipore Sigma)
- 200 nm 680 nm microspheres (F8807, Thermofisher)
- 1M Tris-HCl buffer (15568025, Thermofisher)
- NaCl (BDH8014-2.5K GR, VWR)
- Ultrapure distilled water (10977015, Invitrogen)
- 2-Mercaptoethylamine Hydrochloride (A14377, Alfa Aesar)
- glucose oxidase (G2133, Sigma-aldrich)
- catalase (C30, Sigma-aldrich)
- D-glucose (A16828.36, Thermo scientific)
- 1M Potassium hydroxide (484016, Millipore Sigma)
- 4% Paraformaldehyde (J19943-K2, Thermo scientific)
- 25% Glutaraldehyde(G6257, Sigma Aldrich)
- PBS (J61196.AP, Thermo scientific)
- Triton X 100 (X100, Sigma Aldrich)
- Normal goat serum (50062Z, thermofisher)
- Bovine serum albumin (9048-46-8, Fisher Bioreagents)
- NaHCO_3_ (424270250, Thermofisher)
- Mouse anti-PMP70 antibody (SAB4200181, Sigma Aldrich)
- Donkey anti-mouse antibody (715-005-151, Jackson ImmunoResearch)
- DY634 NHS-ester (634-01, Dyomics)
- rabbit anti-TOM20 antibody (HPA011562, Millipore Sigma)
- Goat anti-rabbit AF 647 (A-21245, Invitrogen)
- Chicken anti-vimentin antibody (AB5733, Millipore Sigma)
- Donkey anti-chicken antibody (703-005-155, Jackson ImmunoResearch)
- CF 660C SE/TFP Ester (92137, Biotium)
- Rat anti-alpha tubulin antibody (MAB1864, Millipore Sigma)
- Donkey anti-rat antibody (712-005-153, Jackson ImmunoResearch)
- Dylight 680 NHS-ester (46419, Thermo Scientific) Nanohole array fabrication equipment (optional)
- ATC Orion Sputter System (AJA International Inc)
- FIB mill (JIB-4700F FIB-SEM, JEOL Ltd.)
- 1x glass slide (12-550-109, Fisher Scientific) Resolution targets
- Negative 1951 USAF Wheel Pattern Test, 3” x 1” (R3L1S4N, Thorlabs)

## Supporting information

Supplemental procedures and Figures

## Acknowledgments and Funding Information

Wei-Hong Yeo thanks the Christina Enroth-Cugell and David Cugell Fellowship for Visual Neuroscience and Biomedical Engineering for their support. We would like to thank Mike Vilches for his assistance with providing images of the Nikon TI2 Solidworks files.

## Disclosures

Cheng Sun and Hao F. Zhang have financial interests in Opticent Inc., which did not support this work. Benjamin Brenner, Wei-Hong Yeo, Youngseop Lee, and Junghun Kweon declare no conflict of interest.

## Code availability

All codes used in this manuscript are available at https://github.com/FOIL-NU/sSMLM-protocol.

## Notes

https://github.com/FOIL-NU/sSMLM-protocol

### Summary of Updates

Supplementary file failed to appear with the previous revision

