## Supplemental procedures and Figures for "Implementation and calibration of spectroscopic single-molecule localization microscopy"

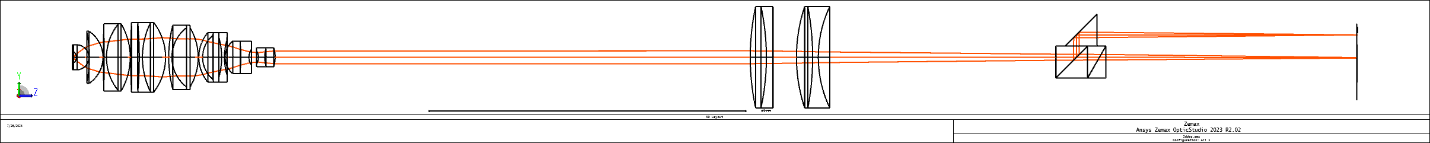

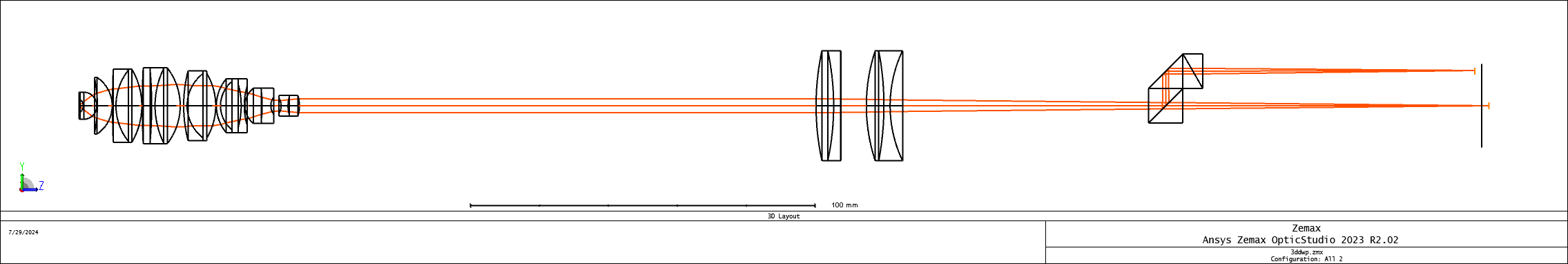

**Figure S1|2D (top) and 3D (bottom) DWP sSMLM system optical detection path comparison**.

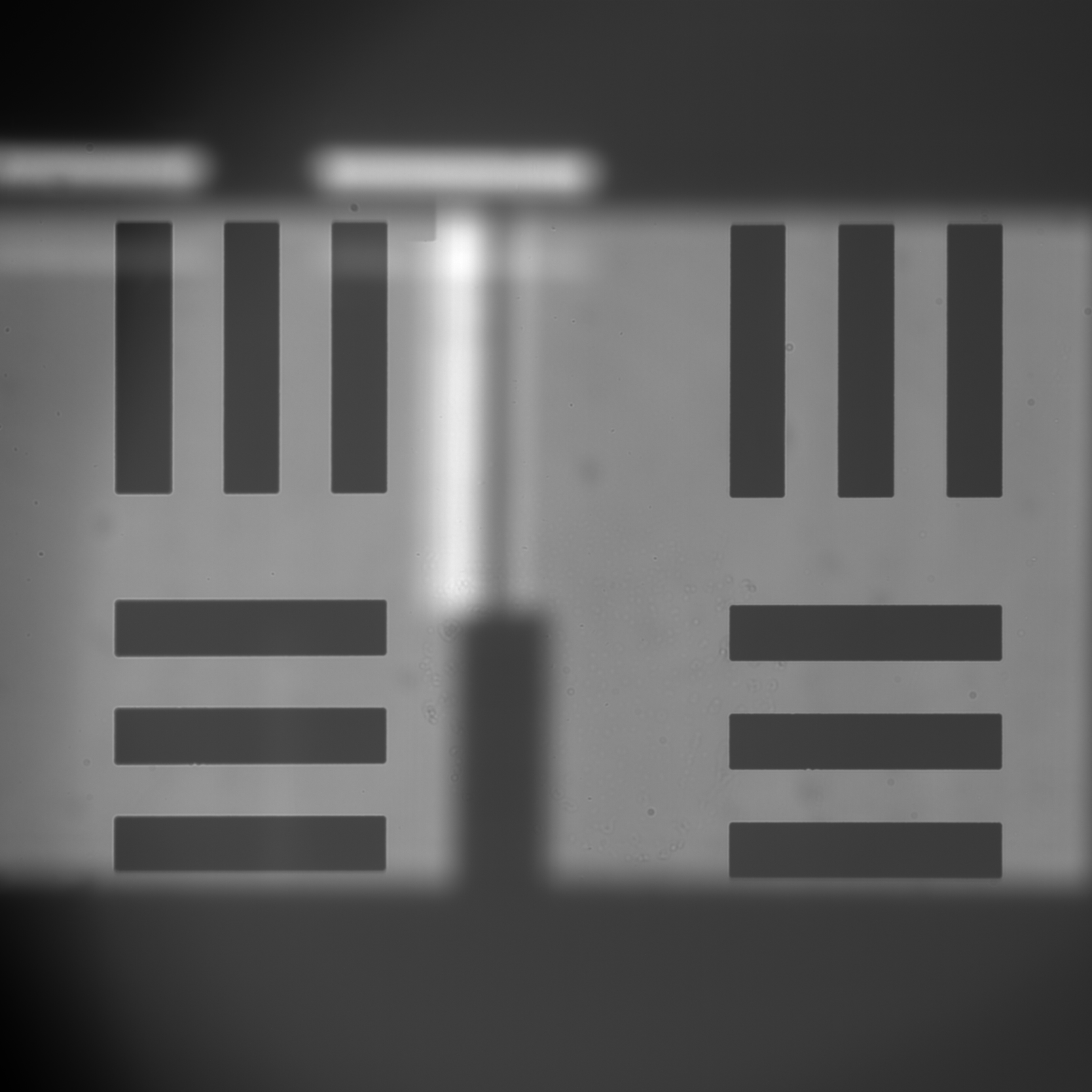

**Figure S2|Sample image of USAF resolution target with the 3D DWP attachment inserted used for spectral calibration.**

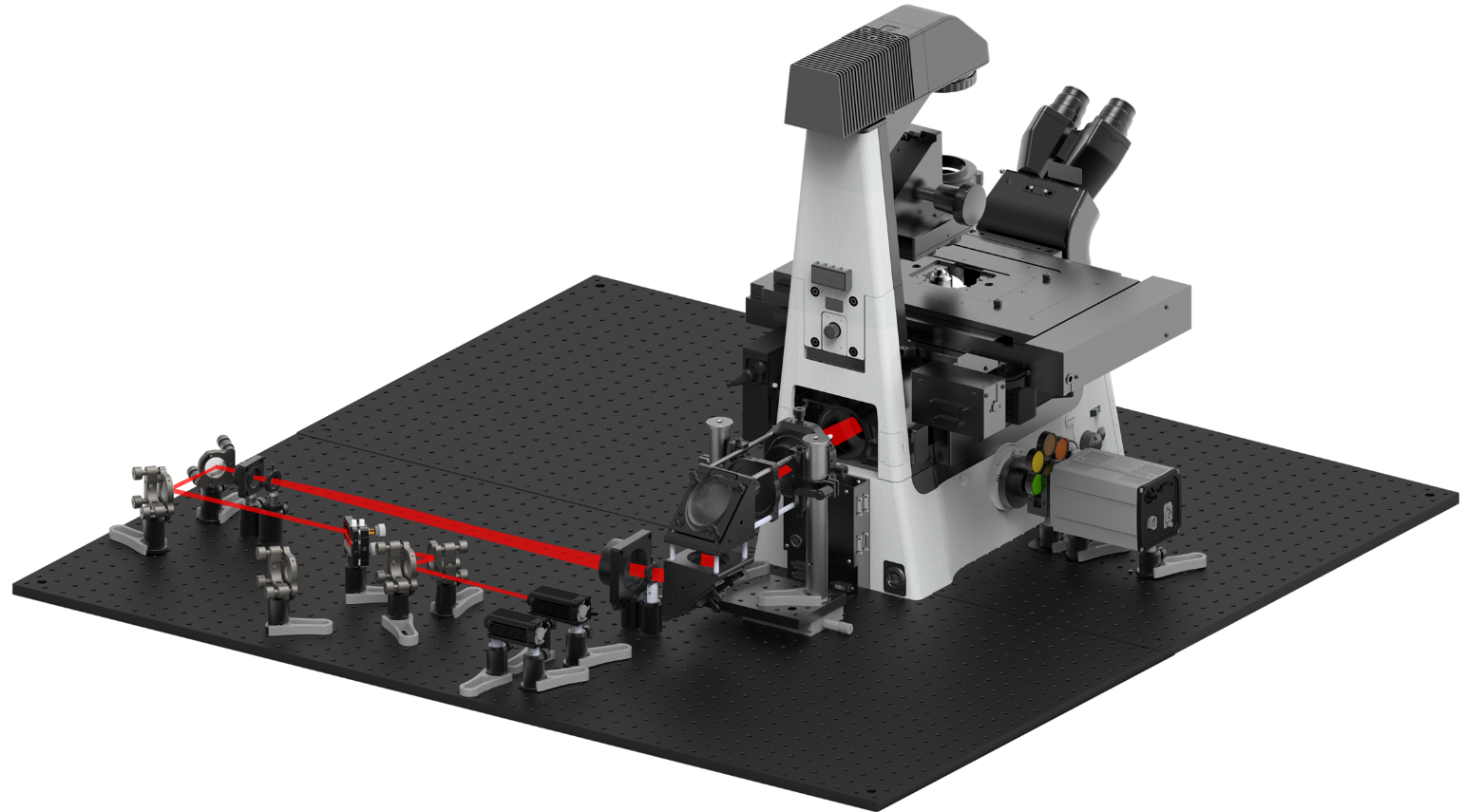

**Figure S3|Full 3D DWP system.**

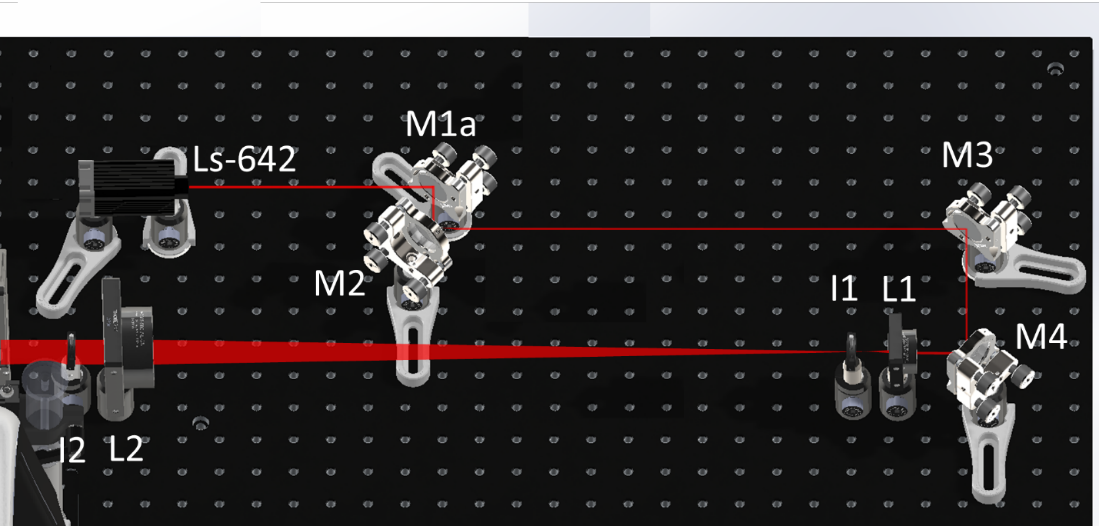

**Figure S4|3D model of the 642 nm laser excitation path.** I: iris.

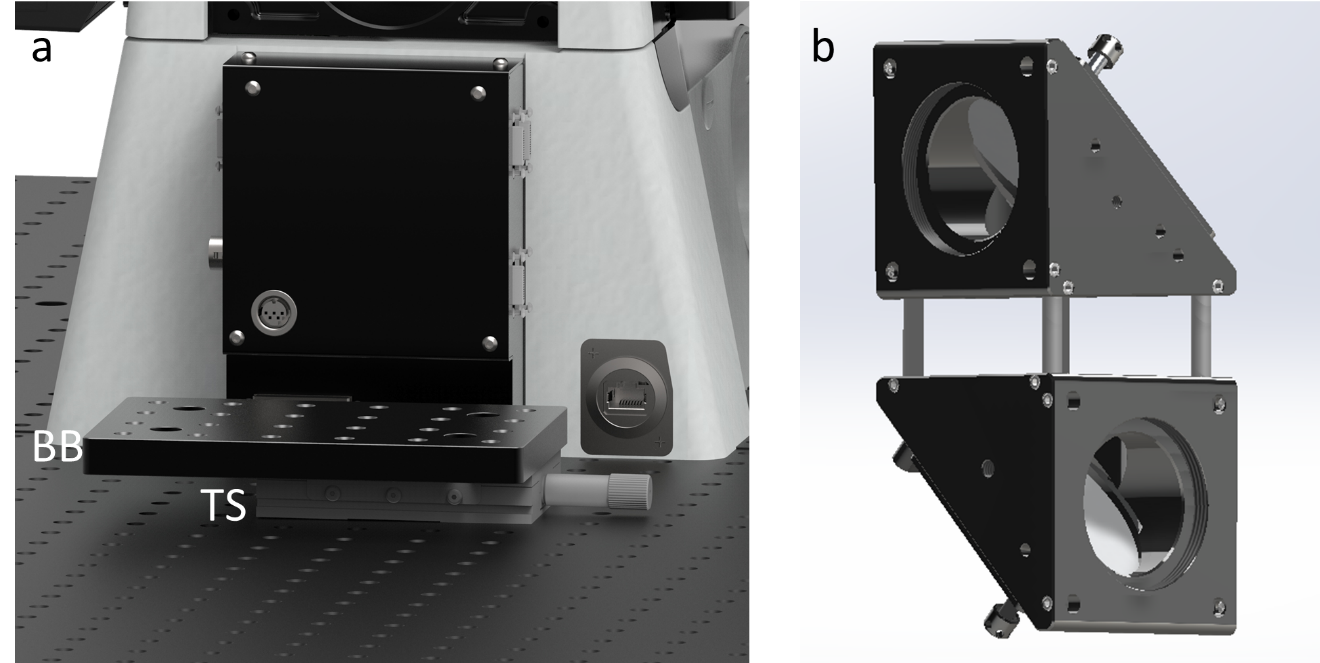

**Figure S5|3D model of the periscope assembly components:** (a) Translational stage with a mounted breadboard behind the microscope body back port. (b) Two elliptical mirrors in mounts set up for the periscope system.

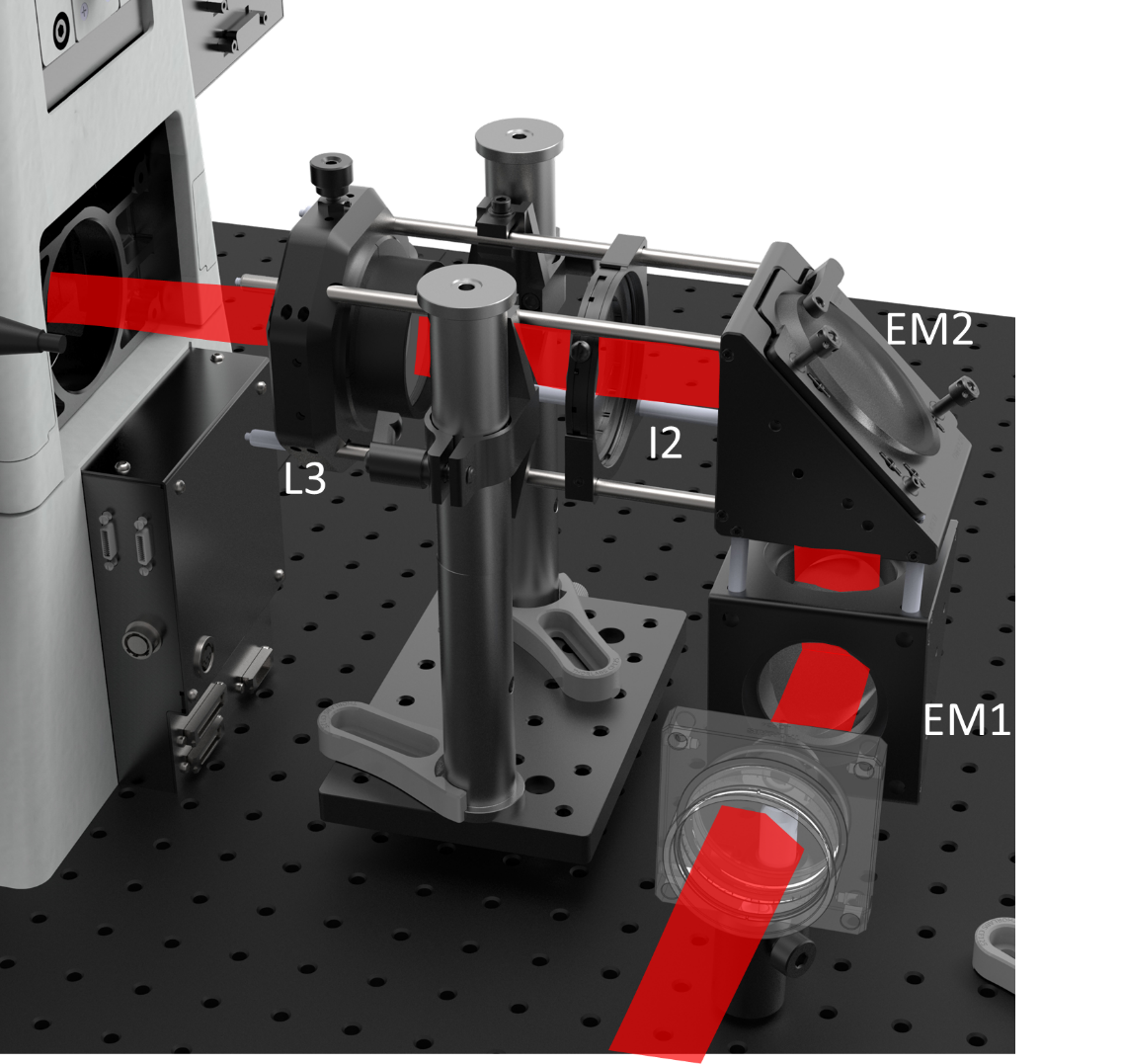

**Figure S6|3D model of the completed periscope system with a superimposed laser line** EM: Elliptical mirror.

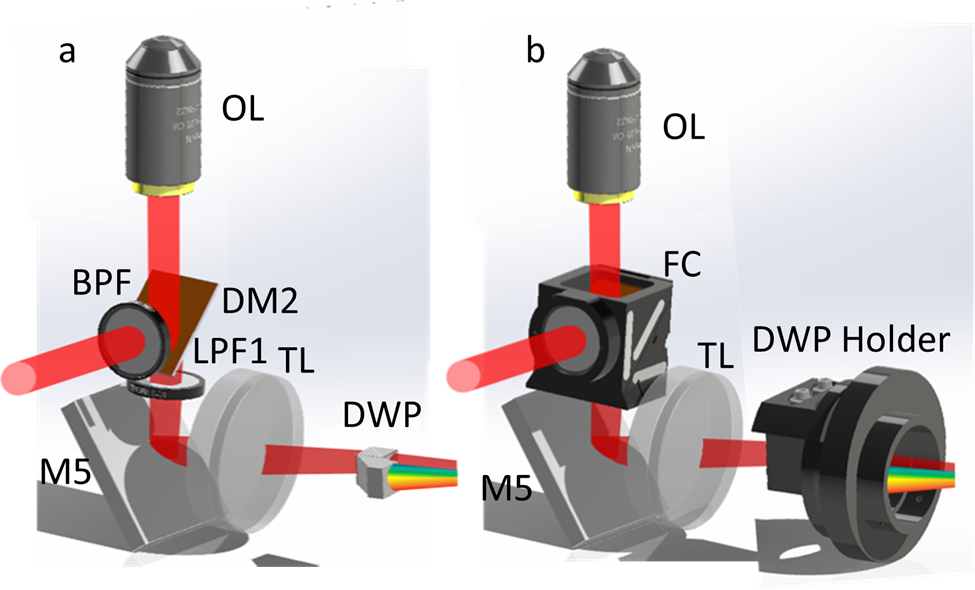

**Figure S7|3D model of components on or within the microscope body with a super-imposed laser line** (a) Optics components without cases. (b) Optics components shown in their cases. FC: Filter cube.

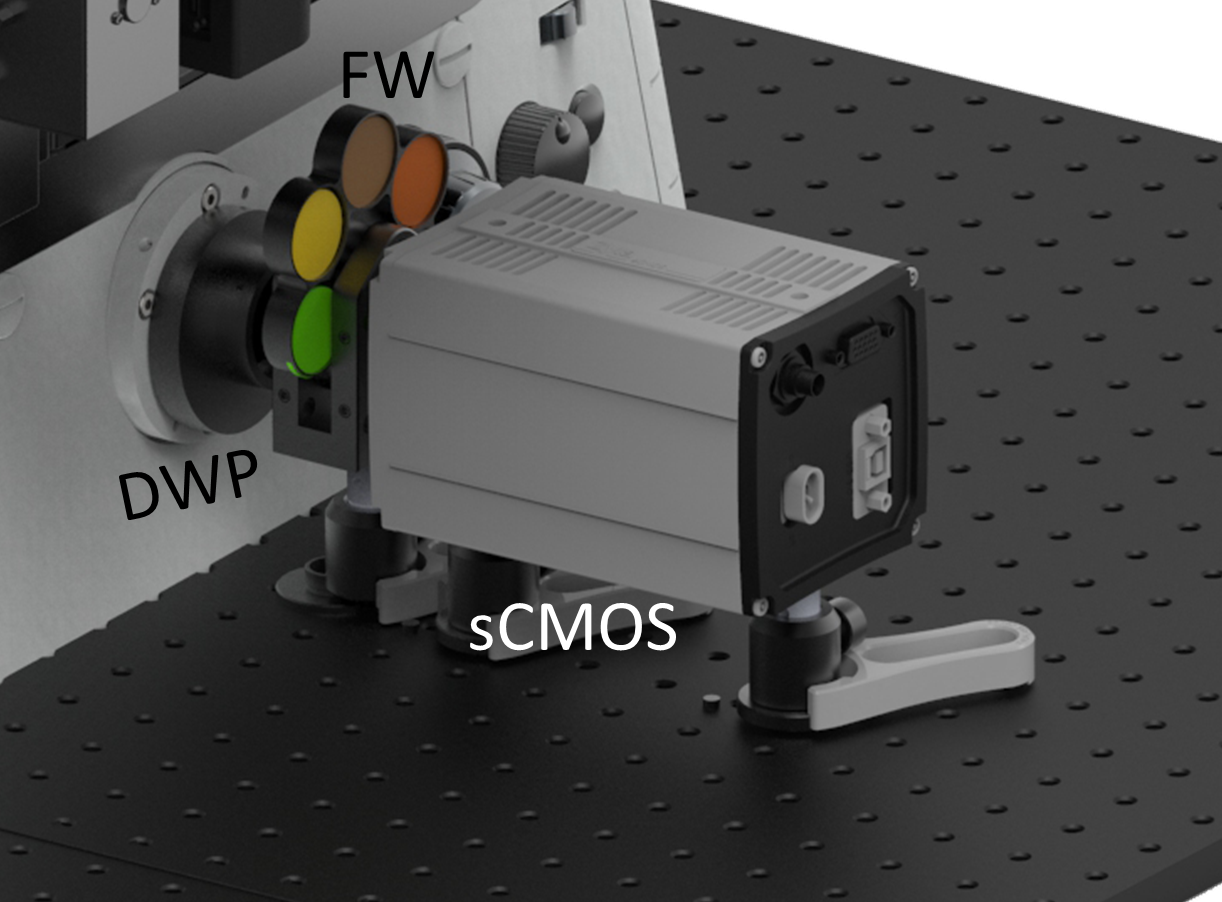

**Figure S8|3D model of the system detection path.**
